# Cell-stereotyped DNA repair outcomes are widespread during genome editing

**DOI:** 10.1101/2025.10.23.684114

**Authors:** Moritz F. Schlapansky, Markus S. Schröder, Antonio J. Santinha, Tanja Rothgangl, Eleonora I. Ioannidi, Grégoire Cullot, Bohdan Lewkow, German Camargo Ortega, Ahmed Nouraiz, Dominic Mailänder, Louisa Selbert, Simon Lam, Aitana Egea, Jiyung J. Shin, Matteo Bordi, Mark DeWitt, Ana Gvozdenovic, Stephen P. Jackson, Timm Schroeder, Helmuth Gehart, Gerald Schwank, Randall J. Platt, Jacob E. Corn

## Abstract

Genome editing outcomes are governed by DNA repair pathways that vary with cell type and state. We developed scOUT-seq (single-cell Outcomes Using Transcript sequencing), a scalable approach that jointly profiles transcriptomes and matched multi-allelic editing outcomes ranging from homology directed repair (HDR) to inter-chromosomal translocations. We mapped editing outcomes in human CD34⁺ hematopoietic stem and progenitor cells (HSPCs), mouse LSK HSPC equivalents, human upper airway organoids, and mouse multi-organ *in vivo* editing. Profiling 500,000 alleles across 74 cell types, scOUT-seq revealed that outcomes in most cell subtypes differ markedly from the bulk average. Various cell types shifted major repair classes, preferred different molecular sequences, and even enriched large structural variants, with distinctive patterns of allelic co-occurrence. Surprisingly, rare stem subtypes diverged from prevalent progenitors, and inhibitory neuron subtypes efficiently incorporated HDR alleles. These data suggest the potential for tailored therapeutic editing that may have been missed by bulk measurements.

## Introduction

The DNA damage response (DDR) continuously monitors the genome, senses diverse forms of damage, and coordinates signaling cascades that enlist cell-cycle checkpoints, chromatin remodeling, and repair machinery^1^. Among the lesions that trigger the DDR, DNA double-stranded breaks (DSBs) occur in virtually all cells of living organisms, both as part of physiological processes such as DNA metabolism or meiotic recombination and as a result of exposure to chemicals or radiation^2^. DSBs are among the most deleterious DNA lesions, since separation from a centromere potentially leads to loss of large amounts of genomic information during mitosis^3^. Repairing DSBs is hence highly conserved, shared across species as diverse as humans and yeast, and the pathways that safeguard genomic integrity are essential for cellular viability and organismal survival^2^.

Despite the deep conservation of core DNA repair pathways, the DDR is subject to extensive metazoan cell specialization. Different cell types of multicellular organisms are exposed to different DNA damaging stresses and express distinct DNA repair factors^1,4^. Genetic diseases linked to mutations in DSB repair genes often exhibit tissue-specific consequences, ranging from cancer to premature aging to neurodegeneration^5–7^. Innovative technologies to map the locations of DNA damage at single cell resolution have revealed that various cell types are prone to DNA damage at particular genomic sites. For example, neurons accumulate DSBs at promoters of ≤1 kb, 5′-untranslated regions and gene bodies^8^. Even within a single tissue, cell types differ sharply in their DNA damage landscapes and repair pathway choice can depend on cell type^9,10^. Chromatin state and gene expression plays a role in both lesion accrual and repair, potentially driving differences between different loci or at a single locus with variable expression^11–13^. However, much less is known about the molecular outcomes resulting from repair at otherwise identical sites in different cells. This aspect of DNA repair is particularly salient in light of the rapid growth of genome editing technologies.

The same cellular machinery responsible for maintaining genome stability is harnessed by genome editing technologies for research and therapeutic purposes^14^. These technologies, most recently exemplified by CRISPR-Cas editors, enable targeted genetic modifications at scale^15^. Although editing technologies differ in their mechanisms of action, the outcome of each edit is ultimately shaped by the cell’s specific DNA repair response. The sequence and frequency of various editing outcomes resulting from DSBs induced by Cas9 are reproducible within a given cell type, non-random and at least partially predictable using machine learning algorithms^16,17^. It has even been suggested that the editing outcome at a particular locus may be static, such that repeated edits in different cells could yield the same outcome each time^18^.

However, functional genomics studies have revealed cell cycle and DNA repair factors that shape the outcomes of CRISPR-Cas mediated editing in human immortalized cells^19–22^. Knockdown of numerous genes can influence editing outcomes, including Fanconi anemia (FA) pathway genes involved in homology directed repair (HDR), POLQ associated with microhomology-mediated end joining (MMEJ), and Pol-λ linked to specific classes of small insertions^19–21,23^. These findings emphasize that genome editing outcomes are intricately tied to genetic background and imply that the expression profiles of particular cell types might shape their editing outcomes. Single-cell genotyping following DSB-based genome editing of approximately 10,000 primary human T cells has recently revealed that even unperturbed cells can acquire a unique constellation of indels and structural variants^24^. However, without matched transcriptomes, it has so far not been possible to determine which cell subtypes favored each editing pattern. To date, there are no large-scale maps linking heterogenous “normal” cells with distinct expression profiles to variable editing outcomes.

To enable the study of tissue- and cell-type-specific editing outcomes, we developed single-cell Outcome using Transcript-sequencing (scOUT-seq). scOUT-seq simultaneously measures transcriptomes and molecular editing outcomes at single-cell resolution. By design, scOUT-seq operates at any scale accessible to standard polyA-capture single cell RNA-seq technologies and in any cell or tissue accessible to these methods. Using scOUT-seq, we mapped the molecular outcomes of genome editing in several heterogeneous contexts: CD34^+^ human hematopoietic stem and progenitor cells (HSPCs), mouse LSK HSPC equivalents, human upper airway organoids, and multi-organ *in vivo* editing of a living mouse. These maps of differential primary cell editing outcomes comprise 9 tissues, 1.2 million cells across 74 cell types, and approximately 500,000 individual alleles. These data are available in browsable form at [XYZ].

We find that, as a rule, editing outcomes vary greatly yet stereotypically between cell subtypes of a given tissue. The bulk repair outcomes measured by standard next-generation sequencing (NGS) is dominated by the most prevalent cells of that tissue, overwhelming contributions from biologically relevant minorities such as stem cells. The differences in editing outcome are not subtle, with various cell subtypes entirely shifting to insertions versus deletions, favoring particular molecular sequences, and even over-representing large structural variants (SVs). These data have therapeutic implications, as for example the rare stem cells within a primary cell mixture can harbor very different outcomes than more prevalent differentiated cells. scOUT-seq and the resulting data have broad application to help guide genome editing for research and therapy and could in future yield rational models for how complex genomic outcomes are shaped by cellular programs.

## Results

### Accurate measurement of molecular editing outcomes and cell types with scOUT-seq

We sought to develop a method that links allelic editing outcomes to single-cell transcriptomes in any cell or tissue type of interest using widely available technology. We started from Genotyping of Transcriptomes, which uses targeted amplification of mRNA transcripts as a proxy for genomic DNA sequence^25^. While Genotyping of Transcriptomes and related technologies are designed to measure a user-specified single nucleotide polymorphism within a gene of interest expressed in a subset of cells, scOUT-seq and its associated bioinformatics pipeline scOUT-MAP (scOUT Mapping and Analaysis Pipeline) extends this capability to detect and classify all potential DNA editing outcomes, including insertions, deletions, HDR, and large-scale structural variants in all the cells of a complex mixture at ∼1,000x the processing speed of other methods (**Figure S1A**)^25,26^.

The general scOUT-seq approach introduces a genome edit into the 3’ untranslated region (UTR) of a carefully selected and ubiquitously transcribed “sentinel” gene (**Figure 1A**). For mouse cells, we tested 55 sgRNAs targeting the 3’ UTR of ubiquitously expressed sentinel genes (e.g. *Eef2*, *Actb*, *Tubb5*, *B2m*, *Psmd4*, *Eif3b*). We eventually selected sgRNAs targeting *Gpi1, Pgk1, B2m*, and *Son* due to their high editing efficiencies, diverse editing profiles, high expression levels of the sentinel gene across multiple cell types, and minimal effects of editing upon the expression level of the sentinel (**Figure S1B-D**). We optimized primer design for each sentinel to efficiently amplify a large region starting upstream of the editing site. By placing one primer within the constant read1 handle, we amplify even structural variants where one side of the edited allele is no longer nearby the original allele (**Figure 1A**). We also included an HDR template (e.g. a single-stranded oligodeoxynucleotide, ssODN) to introduce HDR alleles. After preparing the cell-barcoded polyA single transcriptome library, targeted amplification of a portion of the RNA yields an editing outcome cDNA library. We developed a computational strategy called scOUT-MAP that incorporates enhanced barcode salvaging, an efficient alignment strategy with detailed outcome classification, and explicit linkage between read sequences and alignments, which enables scalable analysis of large combinatorial indexing–based scRNA-seq datasets^27,28^. The alleles contained within each cell are inferred from UMI-normalized read count percentages, flexibly reconstructing the multi-allelic editing outcome. Editing outcomes are represented in a variety of granularities, ranging from broad classifications (e.g. non-homologous end joining (NHEJ) vs HDR) to detailed categories (e.g. PAM-proximal 1-bp insertions) to fully molecular sequences (see Methods).

**Figure 1.**
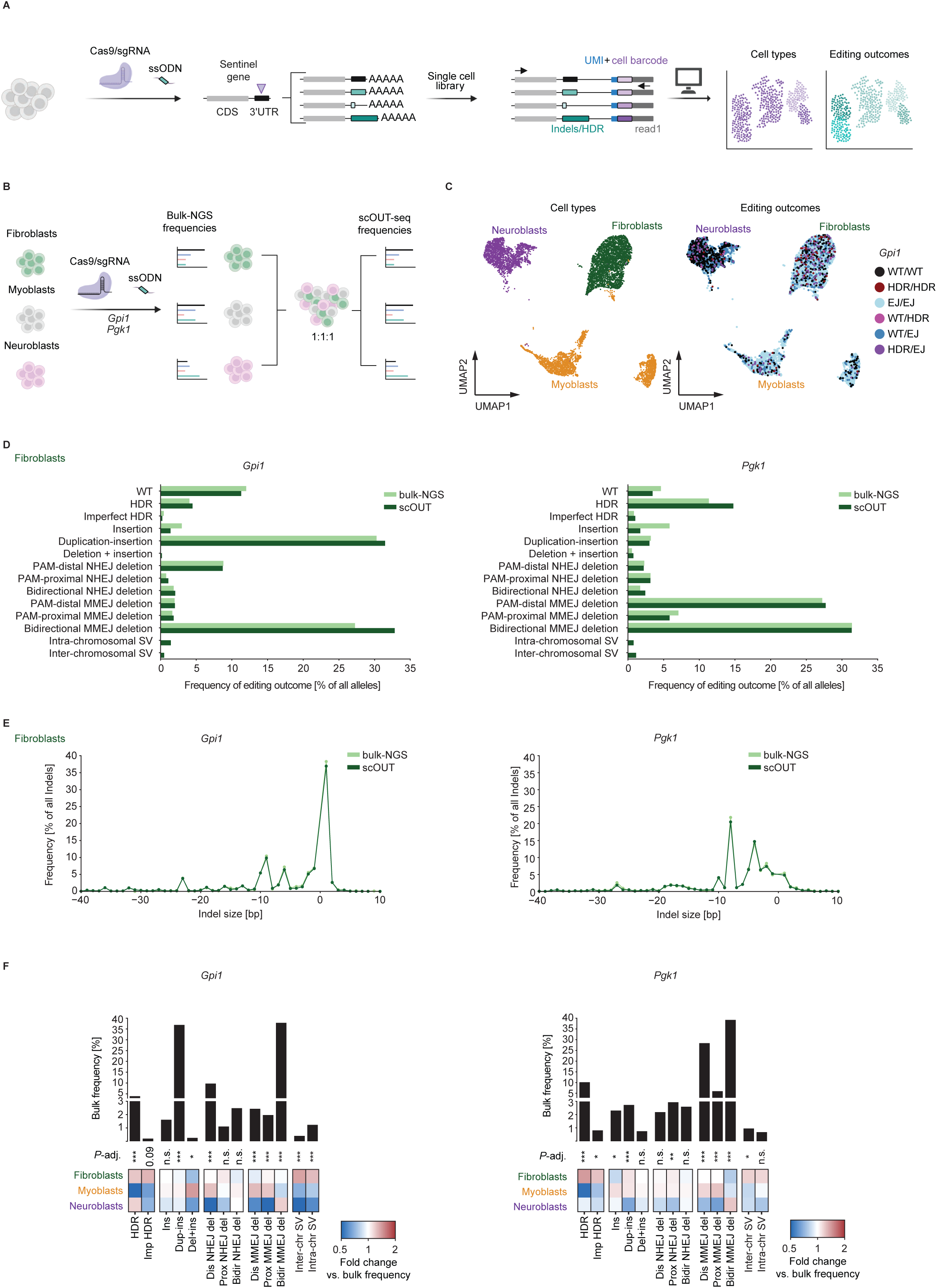
scOUT-seq accurately reflects bulk editing profiles. (A) scOUT-seq workflow. Cas9 ribonucleoproteins (RNPs) and a single stranded oligodeoxynucleotide (ssODN) are used to target 3’ untranslated regions (UTRs) of the sentinel gene which is constitutively and highly expressed across cell types. Cell identity and repair outcomes are then assigned using single-cell sequencing and an associated bioinformatics pipeline (scOUT-MAP). (B) Schematic representation of scOUT-seq testing on defined mixtures of immortalized mouse cell lines. (C) UMAP plots displaying 7,193 cell identities (*left*, color coded by cell line) and editing outcome per cell in 7,043 cells (*right*, color coded by editing outcome at *Gpi1*). WT, wild type; HDR, homology-directed repair; EJ, end-joining. (D) Editing outcomes frequencies at *Gpi1* (*left*) and *Pgk1* (*right*) loci in NIH-3T3 mouse fibroblast cells determined by bulk-NGS and scOUT-seq. WT, wild type (unedited); PAM, protospacer-adjacent motif; NHEJ, non-homologous end-joining; MMEJ, microhomology-mediated end-joining; SV, structural variant. (E) Indel-size frequencies at *Gpi1* (*left*) and *Pgk1* (*right*) loci in NIH-3T3 determined by bulk-NGS and scOUT-seq. bp, base pair. (F) Bar graph displaying bulk frequencies of editing outcome categories in mouse myoblast, fibroblast and neuroblast cell lines (*top*). Hierarchically clustered heatmap of cell-line-specific fold-changes relative to bulk frequencies (*bottom*). FDR-adjusted p-values of G-tests determining difference of categories across cell lines are shown (*, *p*-adj. < 0.05; **, *p*-adj. < 0.01; ***, *p*-adj. < 0.001). Data for *Gpi1* (*left*) and *Pgk1* (*right*) are shown. See also **Figure S1 and S2**.

We first tested scOUT-seq using defined mixtures of immortalized mouse cells. We used a Cas9 ribonucleoprotein (RNP) in combination with HDR ssODNs to separately introduce a DSB in the 3′ UTRs of *Pgk1* or *Gpi1* in individually cultured NIH-3T3 fibroblast cells, Neuro-2a neuroblasts and C2C12 myoblast cells. After editing but before sOUT-seq library preparation, we amplified the edited loci and analyzed them by bulk NGS of PCR amplicons (“bulk outcomes”) (**Figure 1B**). Each edited cell line pool was then mixed in an equimolar ratio (1:1:1 fibroblast:neuroblast:myoblast) and scOUT-seq was used to read out the transcriptomes and paired genotypes (**Figure 1C**). We found that we could accurately deconvolute each cell type within the mixture from their single cell RNA transcriptomes (**Figure 1C, S1E**). Using the scOUT-seq computational pipeline, we successfully assigned single-nucleotide resolution editing outcomes to 7,043 out of 7,193 cells (∼98%) at the *Gpi1* target and to 6,674 out of 7,193 cells (∼93%) at the *Pgk1* target (**Figure 1C, S1F, G**). We found that scOUT-seq amplification led to >50X greater detected UMIs per cell and >2,000x detected reads per cell, as compared to directly analyzing scRNA-seq reads at comparable sequencing depth. Overall, scOUT-seq increased the number of cells with a detectable editing outcome by ∼10-100x relative to direct RNA measurement (**Figure S1H**).

We binned each molecular sequence into one of several descriptive outcomes that encompass various types of HDR, NHEJ, and MMEJ based on previous studies ^21,29,22^ (**Figure S1I**). We observed that a portion of reads only partly matched the target locus, but after the DSB site were uniquely mappable to a different segment of the genome. These reads represented structural variants, which we mapped as intra-chromosomal (i.e. target-anchored large deletions, inversions) or inter-chromosomal (i.e. target-anchored translocations) events. The ability to measure these dramatic outcomes highlights the flexibility of scOUT-seq (**Figure 1A**).

We compared the frequencies of each descriptive outcome between bulk PCR amplicons from the genomic DNA of each cell type and scOUT-seq outcomes aggregated across each transcriptome-assigned cell type. In all three cell types, we found excellent agreement in bulk amplicon outcomes and scOUT-seq outcomes (**Figure 1D, S2A, B**). This was true when installing edits into either the 3’ UTR of *Gpi1* or *Pgk1*. Going a step further, we compared the single nucleotide indel profiles measured in the bulk pool to aggregated single cell indel profiles, again finding excellent agreement in all cell lines and at both loci (**Figure 1E, S2C, D**).

As expected, the binned and molecularly precise outcomes differed between edits made at *Gpi1* and *Pgk1*, reflecting their underlying sequence composition. However, within each locus we observed differences in the descriptive and molecular outcomes between the three cell lines (**Figure 1F, S2C, D**). For example, at the *Pgk1* locus a -9 deletion (bidirectional MMEJ deletion) was abundant in neuroblast cells, but less prominent in fibroblast and myoblast lines. These differences were apparent in both the bulk and scOUT-seq measurements (**Figure 1D-F, S2A-D**). The increased frequency of bidirectional MMEJ deletions at both loci in neuroblasts was consistent with the elevated and broader expression of the key MMEJ factor PolQ (**Figure 1F, S2E**). Notably, scOUT-seq was able to detect intra- and inter-chromosomal structural variants from the single-cell dataset, while the paired PCR primers used for amplicon NGS could not amplify these rearrangements. Such structural variants were missing from bulk amplicon NGS but were present at frequencies up to 1.5% in scOUT-seq data (**Figure 1D, S2A, B**), with notable differences between the three cell lines (**Figure 1F**). We noted that the abundance of wild type alleles differed between the fibroblasts, myoblasts, and neuroblasts. However, this could arise from differential efficiency of delivery to each cell type. We therefore removed wildtype alleles from all subsequent quantitative analyses.

Overall, our results indicated that scOUT-seq provides an accurate representation of repair outcomes at single-cell resolution, while also suggesting the existence of differential editing outcomes in immortalized cells of different lineages. We further explored this idea in several primary cell and organismal contexts: human and mouse hematopoietic stem and progenitor cells (HSPCs), human upper airway organoids, and in multiple tissues of *in vivo* edited mice. For each experimental system, we evaluated whether cell types exhibited editing outcome profiles differing from the bulk, the co-occurrence of multi-allelic editing outcomes in each cell type, and biases in the molecular sequence composition of each cell type.

### Differential editing outcomes are widespread in primary cell subpopulations

We first investigated the distribution of editing outcomes in human adult mobilized peripheral blood CD34+ hematopoietic stem and progenitor cells (HSPCs) and their mouse equivalent Lin^−^ Sca-1^+^ c-Kit^+^ (LSK) cells. An sgRNA targeting the *GAPDH* 3’ UTR was selected after assessing 17 sgRNAs across *ACTB*, *GAPDH*, *B2M* and *GPI* UTRs. The selected *GAPDH*-targeting sgRNA showed the best combination of editing efficiency, diverse editing profile, and HDR rate in multiple HPSC donors (**Figure S3A, B**). We also performed sgRNA screens to expand the pool of validated candidate sgRNAs for broad applicability across tissues and cell types (**Figure S3C**).

We edited 4 million CD34^+^ HSPCs within the 3’ UTR of *GAPDH* using an electroporated Cas9 RNP and an ssODN HDR template (**Figure 2A**). Two days after editing, we sorted out CD34^+^ CD38^-^ engraftment-enriched cells to increase the proportion of HSCs in the mixture^30^, mixed them 1:1 with sorted CD34^+^ CD38^+^ progenitor cells, and performed scOUT-seq on 15,980 cells using droplet-based single cell barcoding. We successfully assigned cell types to all cells and editing outcomes for 14,660 of them (>90%) (**Figure 2B**). 8,042 of the outcome-assigned cells contained at least one edited allele.

**Figure 2.**
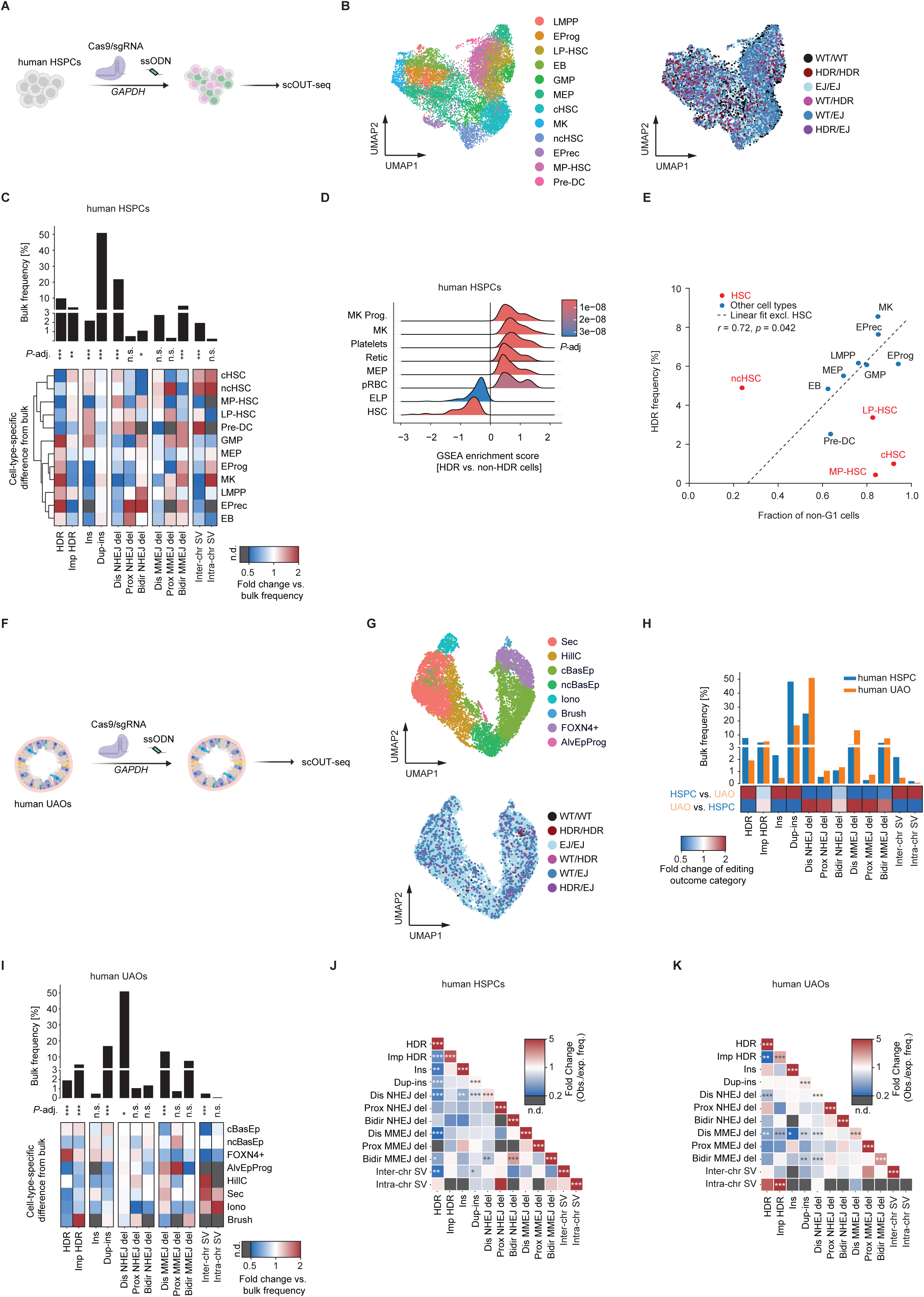
Editing outcome categories in human CD34^+^ HSPCs and upper airway epithelial organoids. (A) Schematic of editing in human CD34^+^ hematopoietic stem and progenitor cells (HSPCs). (B) UMAPs displaying cell identities of 15,980 CD34^+^ HSPCs (*left*, color coded for cell types) and editing outcome of per cell for 14,660 cells (*right,* color coded for editing outcome). LMPP, lympho-myeloid primed progenitors; EProg, erythroid progenitors; LP-HSC, lymphoid-primed HSCs; EB, erythroblasts; GMP, granulocyte-monocyte progenitors; MEP, megakaryocyte-erythroid progenitors; cHSC, cycling HSCs; MK, megakaryocytes; ncHSC; non-cycling HSCs; EPrec, erythroid precursors; MP-HSC, myeloid-primed HSCs; Pre-DC, pre-dendritic cells; WT, wild type; HDR, homology-directed repair; EJ, end-joining. (C) Bar graph displaying bulk frequencies of editing outcome categories in human CD34^+^ HSPCs (*top*). Hierarchically clustered heatmap of cell-type-specific fold-changes relative to bulk frequencies. n.d., not detected (*bottom*). FDR-adjusted p-values of G-tests determining difference of categories across cell lines are shown (*, *p*-adj. < 0.05; **, *p*-adj. < 0.01; ***, *p*-adj. < 0.001; n.s., p-adj. > 0.05). (D) Gene set enrichment analysis (GSEA) comparing HDR versus non-HDR cells using cell-type-specific gene signatures. Distributions are color-coded by FDR-adjusted p-values. MK Prog, megakaryocyte progenitors; MK, megakaryocytes; Retic, reticulocytes, MEP, megakaryocyte-erythroid progenitors; pRBC, primary red blood cells; ELP, early lymphoid progenitors; HSC, hematopoietic stem cells. (E) Scatter plot showing the fraction of non-G1 cells versus HDR frequency across cell types. Linear model excluding HSCs is indicated by the dashed line. r, Pearson correlation coefficient; p, p-value. (F) Schematic of editing in human upper airway organoids (UAOs). (G) UMAP displaying cell identities of UAOs (*top*, color coded for cell types) and editing outcome per cell (*bottom*). Sec, secretory cells; HillC, hillock cells; cBasEp, cycling basal epithelial cells; ncBasEp, non-cycling basal epithelial cells; Iono, ionocytes; Brush, brush cells, FOXN4^+^, FOXN4-positive cells; AlvEpProg, alveolar epithelial progenitors. (H) Comparison of bulk average editing outcome frequencies grouped by categories between human HSPCs and UAOs. Bar graph displaying bulk frequencies of editing outcome categories (*top*). Heatmap of cell-type-specific fold-changes relative to bulk frequencies (*bottom*). (I) Bar graph of bulk frequencies of editing outcome categories in human UAOs (*top*). Heatmap of cell-type-specific fold-changes relative to bulk frequencies. n.d., not detected (*bottom*). FDR-adjusted p-values of G-tests determining difference of categories across cell lines are shown (*, *p*-adj. < 0.05; ***, *p*-adj. < 0.001, n.s., p-adj. > 0.05). (J) Correlation plot showing fold-changes between observed and expected frequency (assuming independence of alleles) of co-occurrence of editing outcome categories in human CD34^+^ HSPCs. n.d., not detected. FDR-adjusted p-values of a χ² (chi-square) test determining difference between observed and expected frequencies across cell types are shown (*, *p*-adj. < 0.05; **, *p*-adj. < 0.01; ***, *p*-adj. < 0.001). (K) Correlation plot of fold-changes between observed and expected frequency (assuming independence of alleles) of co-occurrence of editing outcome categories in human upper airway organoids. n.d., not detected. FDR-adjusted p-values of a χ² (chi-square) test determining difference between observed and expected frequencies across cell types are shown (*, *p*-adj. < 0.05; **, *p*-adj. < 0.01; ***, *p*-adj. < 0.001). See also **Figure S3 and S4**.

We also sorted 406,193 LSK cells from five Cas9-C57BL/6 mice and edited them within the 3’ UTR of *B2m* using AAV-6 expressing the *B2m* sgRNA and containing an HDR template **(Fig. S3D**). Three days post infection, we performed scOUT-seq with droplet-based barcoding on 3,290 cells and assigned editing outcomes to 1,497 of them (**Figure S3E**). 1,461 of the outcome-assigned cells contained at least one edited allele.

HSPCs and LSKs comprise a mixture of cycling and quiescent subpopulations, and prior work has indicated that HDR alleles are over-represented in the cycling cells^31–33^. In human cells, scOUT-seq confirmed that HDR alleles were most abundant in MK, EProg, EPrec and GMP cell types (**Figure 2C, S3F, G**). HSC subpopulations (cHSCs, ncHSCs, MP-HSCs, and LP-HSCs), showed fewer “perfect” HDR alleles (integration of the entire HDR cassette) as compared to the bulk population and far less than cycling cells such as erythroid precursors. However, they showed a slight enrichment for “imperfect” HDR, where only a portion of the HDR cassette was integrated. Unbiased gene set enrichment analysis (GSEA) of transcriptomes from cells with perfect HDR alleles vs those with non-HDR alleles further supported depletion of HDR alleles in HSCs (**Figure 2D**). We observed similar stereotyping of HDR outcomes towards cycling progenitors and away from non-cycling stem subpopulations when editing murine LSKs (**Figure S3H, I**).

We generally observed good correlation between active cell cycle and HDR frequency in human HSPCs (**Figure 2E**). However, human non-cycling HSCs deviated from this relationship, exhibiting more HDR alleles than expected given their quiescence. This is in line with multiple reports of therapeutic levels of HDR in engraftment-capable non-cycling HSCs^31,34,35^, and could reflect transient cycling that enables HDR followed by re-quiescence. However, actively cycling HSCs paradoxically exhibited lower levels of HDR (**Figure 2E**). Mouse LSK cells more strictly adhered to the linear relationship between cycling and HDR with no deviation for non-cycling or cycling HSCs (**Figure S3J**), indicating potential species-specific differences.

While HDR is known to vary depending on cell cycle status and thus cell subpopulation, differential molecular outcomes of end joining arising from genome editing are less well described. We were surprised to find that the bulk average descriptive end joining outcomes in human HSPCs were representative of relatively few subpopulations. Most cell types significantly deviated from the mean (**Figure 2C**). Strikingly, clustering human HSPC cell types solely on their differential binned editing outcomes yielded groupings consistent with their lineage (**Figure 2C, S3F**).

Most human HSPC subpopulations were very different from the bulk average in at least one and sometimes many descriptive editing outcomes. Individual subpopulations balanced one another to yield the bulk average editing outcome rather than representing the bulk average on their own. Multiple HSC subpopulations displayed a stronger tendency toward NHEJ-mediated deletions, whereas more differentiated cell types such as MPs, EProgs, and MKs showed a modest enrichment for MMEJ-mediated deletions (**Figure S3G**). We did not observe a clear correlation between MMEJ tendency and expression level or distribution of *POLQ*, a key mediator of the MMEJ pathway (**Figure S3K**). In murine LSKs the MPCs and MKs exhibited the highest and most broadly distributed expression of *Polq*, yet the highest relative levels of MMEJ were observed in non-cycling HSCs (**Figure S3L**).

The overarching patterns of NHEJ- and MMEJ-mediated deletions could be further resolved into striking fine-grained and cell type–specific differences. Multiple HSC subpopulations displayed a stronger tendency toward PAM-distal NHEJ-mediated deletions, whereas more differentiated cell types such MEPs, LMPPs, EPrecs, and EBs favored PAM-proximal and bidirectional NHEJ. PAM-proximal MMEJ deletions were predominantly observed in non-cycling HSCs, whereas EPrecs and EProgs displayed comparatively fewer such events (**Figure 2C**). Murine ncHSCs also showed an enrichment of PAM-proximal MMEJ deletions, pointing to a potential conserved mechanism underlying these outcomes (**Figure S3I**). Strikingly, therapeutically relevant human cycling and non-cycling HSCs, as well as murine non-cycling HSCs, were enriched for target-anchored intra- and inter-chromosomal structural variants (**Figure 2C, S3J**). These editing outcomes are difficult to detect by classical amplicon NGS methods but their presence in engrafting stem cells could be major barriers to therapy.

To assess whether cell subpopulations in tissues other than mesoderm-derived human and mouse HSPCs exhibit distinct editing outcomes, we performed scOUT-seq in epithelium-derived human upper airway organoids **(Figure 2F)**. The organoids were derived from primary healthy human upper airway epithelium and cultured as 3D tissue structures containing both adult stem cells and the differentiated cell types of the native tissue, with the capacity for self-renewal and long-term propagation *in vitro*. We used Cas9 RNP electroporation to edit the 3’ UTR of *GAPDH* in organoid cells and performed scOUT-seq with droplet-based barcoding to yield 9,774 single cell transcriptomes and 9,643 biallelic editing outcomes (**Figure 2G**). Relative to human CD34^+^ HSPCs, upper airway organoids showed higher NHEJ- and MMEJ-deletion rates (specifically PAM-distal and -proximal NHEJ and MMEJ deletions) and fewer perfect HDR alleles (**Figure 2H**).

Much like HSPCs, the repair outcome frequencies of most organoid cell subpopulations significantly varied between cell types and diverged from the bulk average (**Figure 2I, S3N**). cBasEp displayed higher levels of HDR compared to their non-cycling counterparts (ncBasEp). Within the cycling population, FOXN4⁺ cells - on the trajectory toward ciliated-cell differentiation - exhibited particularly high HDR.”. However as in CD34+ HSPCs, the HDR propensity of some cell types was not solely explainable by cell cycle (**Figure S3N**). End-joining events were also unevenly distributed. PAM-proximal MMEJ deletions were enriched in ncBasEps, while PAM-distal MMEJ deletions were more frequent in Sec, HillC and Iono populations (**Figure 2I**). Notably, variation in *POLQ* expression alone did not account for this distribution of end-joining outcomes (**Figure S3O**). cBasEp showed a slight preference for duplicating insertions, while ncBasEp were enriched for non-templated insertions. We found that perfect HDR was particularly prevalent in FOXN4+, but Brush cells instead over-represented imperfect partial conversion HDR. Inter-chromosomal SVs were predominantly detected in specialized epithelial cell types including Iono, Sec and HillC. Together with results in HSPCs, these findings demonstrate that editing outcomes between and within multiple DNA repair pathways are shaped by cell identity in multiple lineages and organisms.

### Edited allele co-occurrence is biased in single cells

Anecdotal reports suggest that genome editing outcomes might be biased for similar alleles in the same cell^36^. For example, Co-Targeting Selection (CTS) uses a selectable marker to enrich for co-occurrence of a non-selectable event^37^. However, to the best of our knowledge this has not been systematically investigated across multiple cell types at the single cell level.

We leveraged the allele-level resolution of scOUT-seq to compare the observed co-occurrence frequency of each editing outcome pair to its expected frequency, assuming independent distribution of the alleles to each cell under a multinomial model. Consistent with prior reports we found that the descriptive binned repair outcomes co-occur more frequently with themselves than expected by chance in human and mouse HSPCs (**Figure 2J, S3P**) and in human upper airway organoids (**Figure 2K**).

However, we also observed several additional co-occurrences and mutual exclusivities between non-identical outcome types. In HSPCs, intrachromosomal SVs were found more often than expected in cells harboring PAM-proximal NHEJ deletion alleles. Bidirectional MMEJ deletions co-occurred significantly more frequently than expected with bidirectional NHEJ deletions, consistent with a shared reliance on end resection (**Figure 2J**). In HSPCs and organoids, both PAM-distal and bidirectional MMEJ deletions were under-represented with duplication-insertions and PAM-distal NHEJ deletions, further suggesting competition between resection-dependent and c-NHEJ-type sub pathways (**Figure 2J, K, S3P**).

We also observed that inter-chromosomal SVs were mutually exclusive with HDR events in human and mouse HSPCs, while intra-chromosomal SVs occurred slightly more often than expected with HDR in human HSPCs, mouse HSPCs, and human UAOs (**Figure 2J, K, S3P).** We found a significant negative correlation between HDR and inter-chromosomal SV rates in almost all human HSPC cell types, and a similar trend was also observed in human UAOs and murine LSKs (**Figure S4A-C**). These measured biallelic preferences indicate that certain co-occurrence tendencies are conserved across tissues, potentially reflecting a shared bias for cell identity or state that remains to be mechanistically described.

### Molecular editing outcomes differ between cell subpopulations

Having observed cell type specific biases in descriptively binned editing outcomes, we next moved a level deeper to investigate the complex molecular details of each editing outcome. Machine learning models have enabled the prediction of molecular editing outcomes at a given sequence^16,17^. These models typically perform best in the same cell context where they were trained, implying that the host cell plays a role in determining the molecular outcome, even within broader bins. We first compared the distribution of indel sizes at single nucleotide resolution in individual cell types to the bulk population average.

In human HSPCs, we found that the bulk indel spectrum was a population-weighted average of the differential molecular outcomes of the component cell types. The indel size distribution of many cell types was not only distinct from the bulk average, but also from one another (**Figure 3A**). MEPs most closely represented the bulk average, but in general no single cell type was identical to the bulk. The differential molecular outcomes included a marked switch in insertions versus deletions between cell types. HSC-related cells (ncHSC, cHSC, LP-HSC, and MP-HSC) exhibited a bias towards smaller deletions, while more differentiated cell types (EPrec, EProg, RBC, and MK) showed an enrichment for larger deletions and single-base insertions (**Figure 3A**). The sole exception was pre-DCs, which displayed an HSC-like indel spectrum. The HSC-related populations from murine LSK cells exhibited a similar bias towards smaller deletions, implying a conserved biology that somehow shapes indel outcomes (**Figure S4D**).

**Figure 3.**
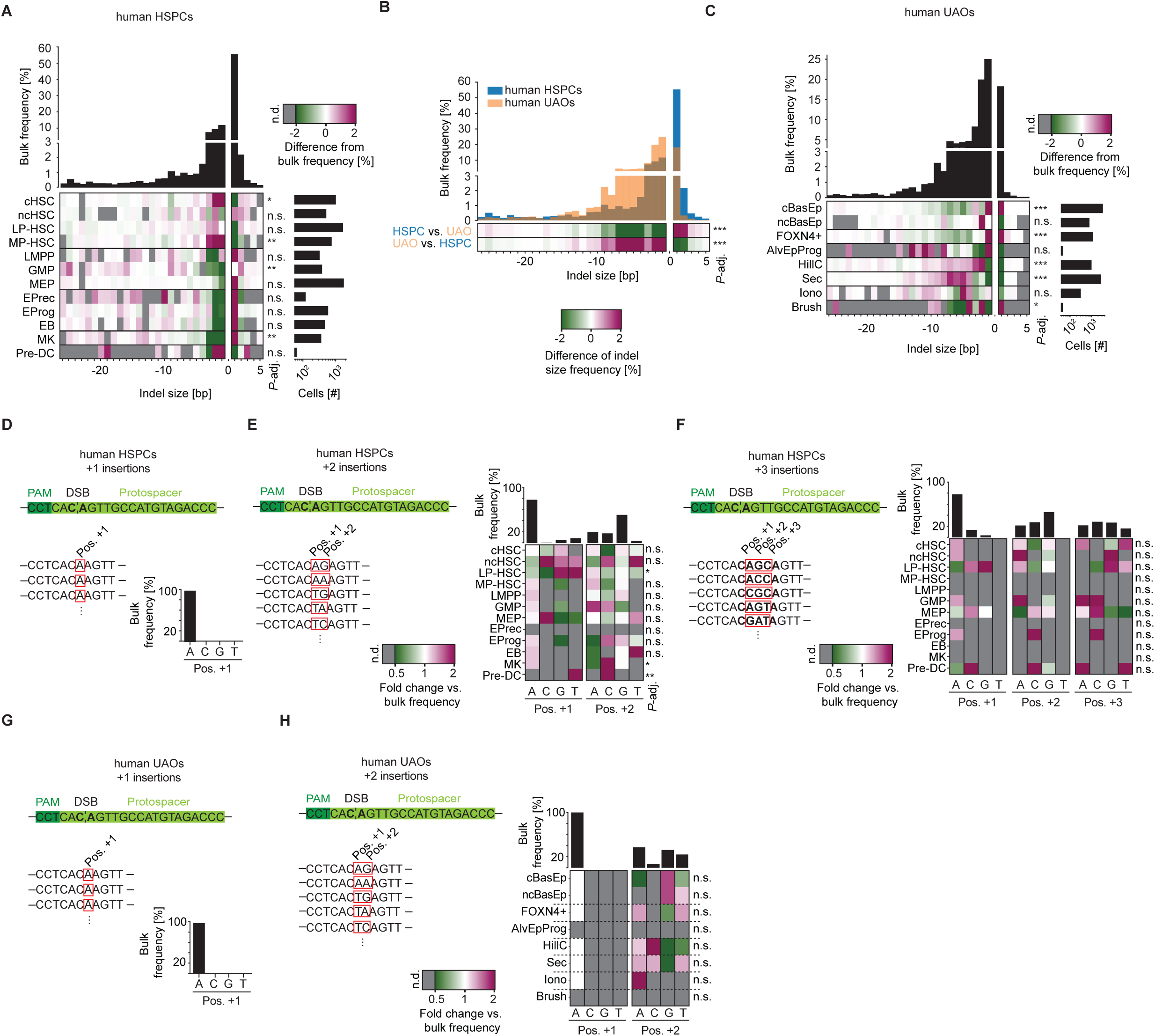
Molecular editing outcomes in human CD34^+^ HSPCs and upper airway epithelial organoids. (A) Histogram showing bulk frequencies of indel sizes in CD34^+^ HSPCs (*top*). Heatmap of percentage-point deviations in indel-size frequencies from the bulk across cell types. n.d., not detected. (*bottom left*). The number of cells per cell type is depicted in a bar graph (*bottom right*). FDR-adjusted p-values of G-tests determining difference of indel profile to bulk are shown (**; *p*-adj. < 0.01; ***; *p*-adj. < 0.001, n.s.). LMPP, lympho-myeloid primed progenitors; EProg, erythroid progenitors; LP-HSC, lymphoid-primed HSCs; EB, erythroblasts; GMP, granulocyte-monocyte progenitors; MEP, megakaryocyte-erythroid progenitors; cHSC, cycling HSCs; MK, megakaryocytes; ncHSC; non-cycling HSCs; EPrec, erythroid precursors; MP-HSC, myeloid-primed HSCs; Pre-DC, pre-dendritic cells. (B) Histogram overlay of bulk indel-size frequencies in human CD34^+^ HSPCs and upper airway organoids (UAOs) (*top*). Heatmap indicating percentage-point deviations in indel-size frequencies between human HSPCs and UAOs (*bottom*). (C) Histogram showing bulk frequencies of indel sizes in human UAOs (*top*). Heatmap of percentage-point deviations in indel-size frequencies from the bulk across cell types. n.d., not detected. (*bottom left*). The number of cells per cell type is depicted in a bar graph (*bottom right*). FDR-adjusted p-values of G-tests determining difference of indel profile to bulk are shown (*, *p*-adj. < 0.05; ***, *p*-adj. < 0.001; n.s., p-adj. > 0.05).). Sec, secretory cells; HillC, hillock cells; cBasEp, cycling basal epithelial cells; ncBasEp, non-cycling basal epithelial cells; Iono, ionocytes; Brush, brush cells, FOXN4^+^, FOXN4-positive cells; AlvEpProg, alveolar epithelial progenitors. (D) Bar graph of bulk sequence composition of +1 insertions in CD34^+^ HSPCs. The sequence context of the *GAPDH* sgRNA, as well as examples of single base pair insertions, are depicted as a reference. (E) Insertion sequence composition of +2 insertions in CD34^+^ HSPCs. Fold change compared to the bulk is plotted in a heatmap across cell types and insertion sequence positions. n.d., not detected. Bar graphs depict the bulk frequency of each base at each position. The sequence context of the *GAPDH* sgRNA, as well as examples of single base pair insertions, are depicted as a reference. FDR-adjusted p-values of G-tests determining difference of insertion sequence composition to bulk are shown (*, *p*-adj. < 0.05; **, *p*-adj. < 0.01; n.s., p-adj. > 0.05). (F) Insertion sequence composition +3 insertions in CD34^+^ HSPCs. Fold change compared to the bulk is plotted in a heatmap across cell types and insertion sequence positions. n.d., not detected. Bar graphs depict the bulk frequency of each base at each position. The sequence context of the *GAPDH* sgRNA, as well as examples of single base pair insertions, are depicted as a reference. FDR-adjusted p-values of G-tests determining difference of insertion sequence composition to bulk are shown (n.s., p-adj. > 0.05). (G) Bar graph of bulk sequence composition of +1 insertion in human UAOs. The sequence context of the *GAPDH* sgRNA, as well as examples of single base pair insertions, are depicted as a reference. (H) Insertion sequence composition of +2 insertions in human UAOs. Fold change compared to the bulk is plotted in a heatmap across cell types and insertion sequence positions. n.d., not detected. Bar graphs depict the bulk frequency of each base at each position. The sequence context of the *GAPDH* sgRNA, as well as examples of single base pair insertions, are depicted as a reference. FDR-adjusted p-values of G-tests determining difference of insertion sequence composition to bulk are shown (n.s., p-adj. > 0.05). See also **Figure S4**.

Upper airway organoids edited at the same locus as HSPCs exhibited a distinct bulk average indel distribution, characterized by a tendency towards more and larger deletions and a reduced insertion rate (**Figure 3B**). Comparing between cell subpopulations within the organoids, we once again found extensive variability of indel distributions between cell types. Strikingly similar to HSPCs, smaller deletions were more common in less differentiated cells (cBasEp, ncBasEp, FOXN4+), regardless of their proliferative state (**Figure 3C**). More differentiated cell types such as Sec, HillC and Iono exhibited a higher frequency of larger deletions. These findings reveal that cell type shapes indel distributions during Cas9 editing and suggests a conserved biases towards smaller deletions in more stem-like cells.

We next investigated the precise sequence composition of the insertions present in each HSPC subpopulation, ranging from +1 to +3 insertions. In erythroid precursors, only single-nucleotide insertions were observed, whereas in MP-HSCs, LMPPs, and EBs and MKs no insertion exceeded two base pairs (**Figure S4E, F**). Prior studies have suggested that +1 insertions are biased towards templating by the PAM-distal base of the DSB.^17,38^ We also observed this templating of single-base insertions in every cell subpopulation (**Figure 3D**). However, +2 and +3 insertions exhibited greater sequence diversity but marked stereotyping between cell types.

The bulk average +2 insertions strongly over-represented A (templated) at position 1 and weakly over-represented G (potentially also templated) at position 2. This bulk average did not represent even distribution of “AG” insertion alleles across all cell types. Instead, cell-specific propensities formed a population-weighted average to yield the bulk (**Figure 3E**). In position one, non-cycling HSCs exhibited less +1A insertion than the bulk average across all cells, which redistributed to more +1C/G/T. LP-HSCs similarly had less +1A, but only over-represented +1G/T relative to the other cell types. In position two, non-cycling HSCs had far more +2T insertions than any other cell type, while the LP-HSCs now exhibited only minor deviations from the bulk average.

Rarer bulk average +3 insertions were primarily A at position one, relatively heterogeneous at position two but completely lacking T and evenly distributed at position three (**Figure 3F**). EProgs had disproportionately less three base pair insertions, while MEPs had more (**Figure S4F**). Each cell type exhibited characteristic deviations from the +3 insertion average that only partially overlapped with that type’s +2 insertion composition preference. For example, +3 insertions in non-cycling HSCs now over-represented A at the first and second positions relative to other cell types, but G at the third position (**Figure 3F**). LP-HSCs by contrast continued to under-represent A at the first position (as in their +2 insertions) but had relatively little stereotyping in the second position and a strong preference for G at the third position.

We tested whether the nucleotide composition of insertion sequences at an identical targeted site differs between cell types in upper airway organoids. Consistent with findings in HSPCs, +1 insertions predominantly consisted of A across all cell types (**Figure 3G**). As for HSPCs, +2 insertions exhibited greater nucleotide diversity and cell type stereotyping (**Figure 3H**). Position one remained A across cell types, while position two varied in a cell type–specific manner. In basal epithelial cells, regardless of mitotic state, position two was predominantly +2G. In FOXN4⁺ cells, position two was more frequently +2A/T. However hillock cells showed a strong preference for C in the second position, while position two in secretory cells tended to be any base except G. Plus +3 insertions were too rare in UAOs to draw conclusions. Overall, these results reveal that single-base insertions are consistent in their composition across cell types and tissues, but multi-base insertions reflect distinct cell type– and tissue-specific sequence biases in Cas9 editing outcomes. Hence, molecular editing outcomes stemming even from a related pathway (e.g. short insertions) can exhibit cell type specific preferences during genome editing. The mechanistic underpinnings of these differences remain to be determined.

### Perturbing NHEJ elicits cell-type specific effects

Genetic and small molecule perturbations can alter the frequency of editing outcomes within a bulk mixture of cells. It is unknown whether such perturbations equally affect all cells in a heterogeneous population. But the tissue-specific etiology of many DNA repair disorders implies that outcomes in particular cells may be differentially affected by perturbation. We investigated the cell-specific effects of perturbing canonical NHEJ through inhibition of DNA-PKcs, which increases Cas9-driven HDR but also causes larger genomic aberrations^39–42^.

We edited human CD34^+^ HSPCs and human upper airway organoids using electroporation of a Cas9 RNP targeting the 3’ UTR of *GAPDH* and ssODN HDR template in the presence or absence of the DNA-PKcs inhibitor AZD7648 (**Figure 4A, B**). scOUT-seq using droplet-based barcoding of both HSPCs and organoids revealed that AZD7648 redirected bulk editing outcomes towards HDR, and this was reflected broadly across almost all cell subtypes (**Figure 4C, D, S5A, B**).

**Figure 4.**
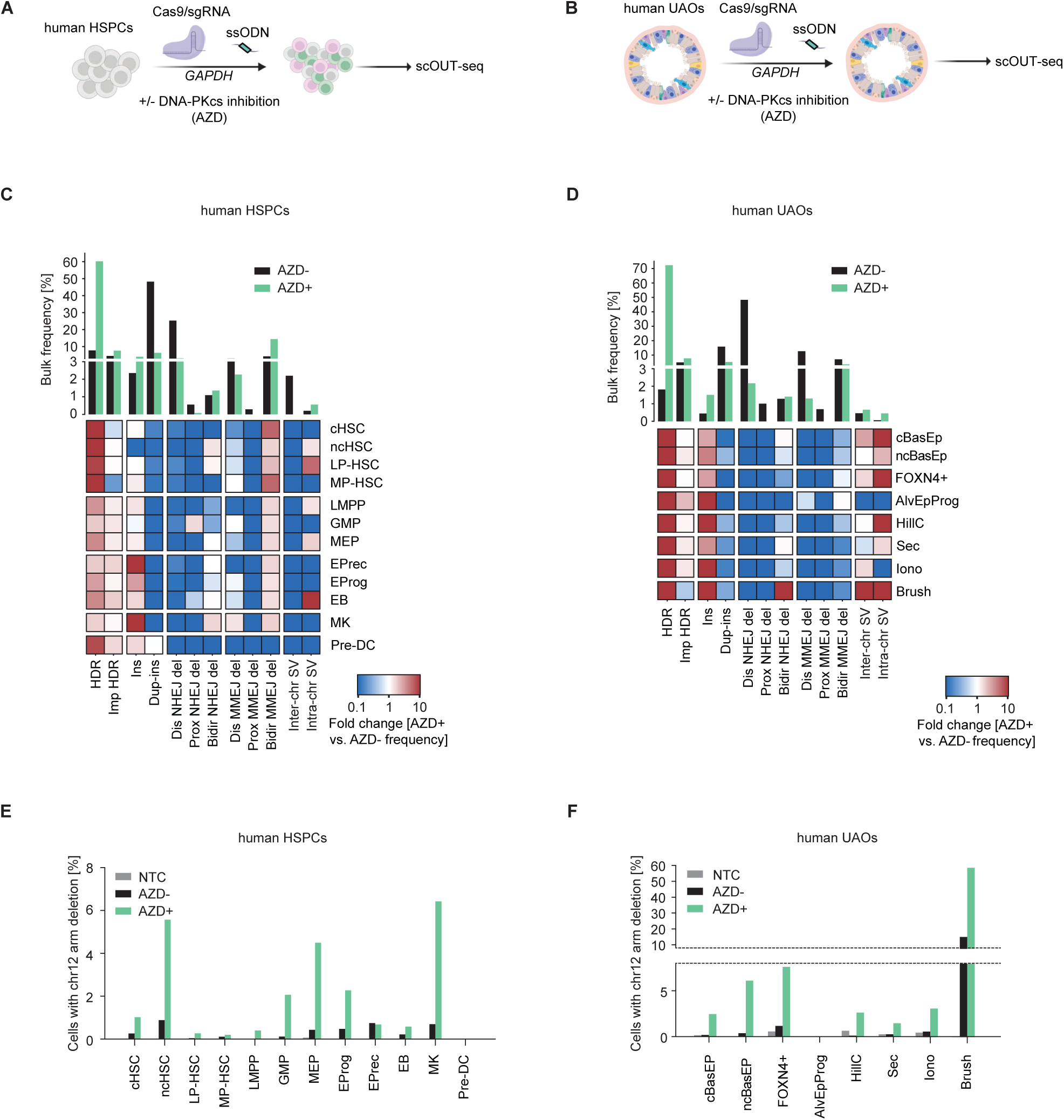
Cell-type-specific effects of DNA-PKcs inhibition during editing in human CD34^+^ HSPCs and upper airway epithelial organoids. (A) Schematic of editing human CD34^+^ HSPCs using DNA-PKcs inhibition (AZD). Cells were treated for 24 hours with 1 µM AZD7648. Untreated but edited cells were used as a control. (B) Schematic of editing human UAOs using DNA-PKcs inhibition. Cells were treated for 24 hours with 1 µM AZD7648. Untreated but edited cells were used as a control. (C) Bar graph showing bulk frequencies of broad editing outcome categories in AZD untreated (AZD-) and AZD treated (AZD+) human HSPCs (*top*). Heatmap of fold-changes in DNA-PKcs inhibited versus non-inhibited CD34*+* HSPCs across different cell types and editing outcome categories (*bottom*). LMPP, lympho-myeloid primed progenitors; EProg, erythroid progenitors; LP-HSC, lymphoid-primed HSCs; EB, erythroblasts; GMP, granulocyte-monocyte progenitors; MEP, megakaryocyte-erythroid progenitors; cHSC, cycling HSCs; MK, megakaryocytes; ncHSC; non-cycling HSCs; EPrec, erythroid precursors; MP-HSC, myeloid-primed HSCs; Pre-DC, pre-dendritic cells; AZD^+^, edited with DNA-PKcs-inhibition; AZD^-^, edited without DNA-PKcs inhibition. (D) Bar graph showing bulk frequencies of broad editing outcome categories in AZD untreated (AZD-) and AZD treated (AZD+) human UAOs (*top*). Heatmap of fold-changes in DNA-PKcs inhibited versus non-inhibited UAOs across different cell types and editing outcome categories (*bottom*). Sec, secretory cells; HillC, hillock cells; cBasEp, cycling basal epithelial cells; ncBasEp, non-cycling basal epithelial cells; Iono, ionocytes; Brush, brush cells, FOXN4+, FOXN4-positive cells; AlvEpProg, alveolar epithelial progenitors. (E) Bar graph depicting the fraction of cells within indicated cell types with chromosome 12 arm deletions across different treatments in CD34^+^ HSPCs. NTC, non-targeting control, AZD+, edited with DNA-PKcs inhibition, AZD-, edited without DNA-PKcs inhibition. (F) Bar graph depicting the fraction of cells within indicated cell types with chromosome 12 arm deletions across different treatments in human UAOs. NTC, non-targeting control, AZD+, edited with DNA-PKcs inhibition, AZD-, edited but untreated. See also **Figure S5 and S6**.

In both HSPCs and organoids, the increase in HDR alleles was accompanied by a decrease in classical NHEJ alleles, consistent with inhibition of DNA-PKcs^43^. Indel spectra broadened in HSPCs upon AZD7648 treatment: larger deletions accumulated at microhomologous sequences (9 and 11 bp), with a corresponding increase in binned bidirectional MMEJ deletions across almost all cell types (**Figure 4C, S5C**). This is consistent with reports that inhibiting DNA-PKcs may divert repair towards *POLQ*-driven alternative end joining^39,41,42^. Larger insertions became more prominent at the cost of 1-bp templated insertions, which binned to an overall increase in non-templated insertions (Ins). However, these non-templated insertions predominantly affected myeloid lineage cells such as EPrecs and MKs. In upper airway organoids, DNA-PKcs inhibition also increased non-templated larger insertions, while decreasing +1 templated insertions (**Figure 4D, S5D**). DNA-PKcs inhibition also decreased bidirectional MMEJ deletions, but only in organoids.

Inhibition of DNA-PKcs reduced target-anchored SVs across all HSPC cell types except EBs but increased these SVs in most cell types in human UAOs (**Figure S5A, B**). When stratifying SVs – as well as other classes – into more detailed categories, we noticed that intra-chromosomal SVs in both HSPCs and organoids were elevated, consistent with prior reports in bulk cells^39,42^ (**Figure 4C).** However, this effect was markedly uneven among cell types. In HSPCs, EBs and LP-HSCs exhibited the greatest increase in intra-chromosomal SVs, while in other cell types these events were unaffected (ncHSC) or even decreased (**Figure 4C, S5E**). UAOs displayed similar heterogeneity: cBasEp, FOXN4+, HillC, and Brush cell types showed pronounced increases in intra-chromosomal SVs, while AlvEpProg and Iono cells showed decreases (**Figure 4D, S5F**). DNA-PKCs inhibition did not simply amplify existing events, as for example LP-HSCs and EBs had fewer intra-chromosomal translocations than the bulk average in untreated samples but increased in the presence of treatment. Conversely, cHSC had more intra-chromosomal translocations in the absence of AZD7648 treatment but decreased in the presence of treatment.

In contrast, target-anchored inter-chromosomal translocations completely disappeared upon AZD7648 treatment in human HSPCs, in line with evidence that these events depend on canonical NHEJ in human cells^44^ (**Figure 4C, S5G**). However, several UAO cell types (cBasEp, FOXN4+, Iono, Brush) exhibited increases in inter-chromosomal SVs, while others showed no change (ncBasEp, HillC) or decreases (AlvEpProg, Sec), indicating cell-specific determinants of target-anchored translocation formation in human cells (**Figure S5H**).

In addition to translocations, inhibiting DNA-PKcs with AZD7648 can cause chromosome arm loss in bulk primary cell populations^39,42^. We inferred chromosome arm loss or gain from single cell expression data across large regions of each chromosome and compared frequencies between cells treated or not with AZD7648^45^. We observed low levels of chromosome arm gain that was mostly unaffected by DNA-PKcs inhibition, but chromosome arm loss was relatively prevalent in the absence of AZD7648 in a cell-type specific manner. (**Figure 4E-F, S6A-C**).

In human HSPCs, editing in the absence of AZD7648 induced detectable chromosome arm deletions extending from the cut site to the telomere of chromosome 12. These deletions were most apparent in ncHSCs, MEPs, erythroid progenitors and precursors, and mature megakaryocytes (**Figure 4E, S6A**). Cell stereotyped chromosome arm losses were also apparent in AAV6-edited murine LSK cells, especially in cHSCs as compared to megakaryocytes and non-cycling HSCs (**Figure S6G-I**). In untreated UAOs, chromosome arm deletions were enriched in ncBasEps, FOXN4+, Iono and Brush cells (**Figure 4F, S6D-F**).

Inhibition of DNA-PKcs dramatically increased chromosome arm loss in HSPCs and UAOs, but the magnitude of this effect differed among cell types rather than affecting them uniformly (**Figure S6C, F**). For example, about 6% of non-cycling HSCs and megakaryocytes lost the arm of chromosome 12 when exposed to AZD7648, whereas MP-HSCs, LMPPs, erythroid precursors and erythroblasts exhibited ≤1% arm loss (**Figure 4E**). Relative fold-change differences in arm loss were widespread, with MP-HSCs, EPregs and EBs exhibiting relatively little change in chromosome arm loss, but GMPs exhibited an over 15-fold increase (**Figure S6C**). Similarly, in UAOs around 6% of ncBasEp cells and 7% of FOXN4+ cells showed chromosome arm loss upon DNA-PKcs inhibition, but only about 3% of cycling basal cells and hillock cells (**Figure 4F**). HillC cells displayed the highest relative increase in chromosome arm loss upon AZD7648 treatment while Sec, Iono and Brush cells were relatively unchanged (**Figure S6F**). Brush cells already harbored substantial baseline arm loss (15 %), which rose to almost 60 % after AZD7648 treatment (**Figure 4F**). However, DNA-PKcs inhibition expanded the Brush cell pool, and we cannot exclude the possibility that chromosome-arm loss promoted brush-cell differentiation (**Figure S6D, F**). Together, these data indicate that perturbations affecting genome editing outcomes can have cell-type specific effects.

### In vivo editing outcomes reveal widespread organ and cell stereotyping

Having found extensive differences in editing outcomes between *ex vivo* edited heterogeneous primary cell sources, we reasoned this might also be reflected during *in vivo* editing of tissues in an intact organism. We focused on using scOUT-seq to query editing in the liver, heart, skeletal muscle, lung and brain, since these organs are major targets for various therapeutic indications and are accessible to AAV-mediated delivery of editing reagents.

We used recombinant AAV vectors targeting multiple tissues to systemically deliver sgRNAs to Cas9-expressing BL6 mice (**Figure 5A**). The AAV vectors contained an sgRNA expression cassette to perform editing and EGFP or dTomato to allow flow cytometry enrichment of infected nuclei. The AAV constructs also included barcoded HDR donors when targeting the brain.

**Figure 5.**
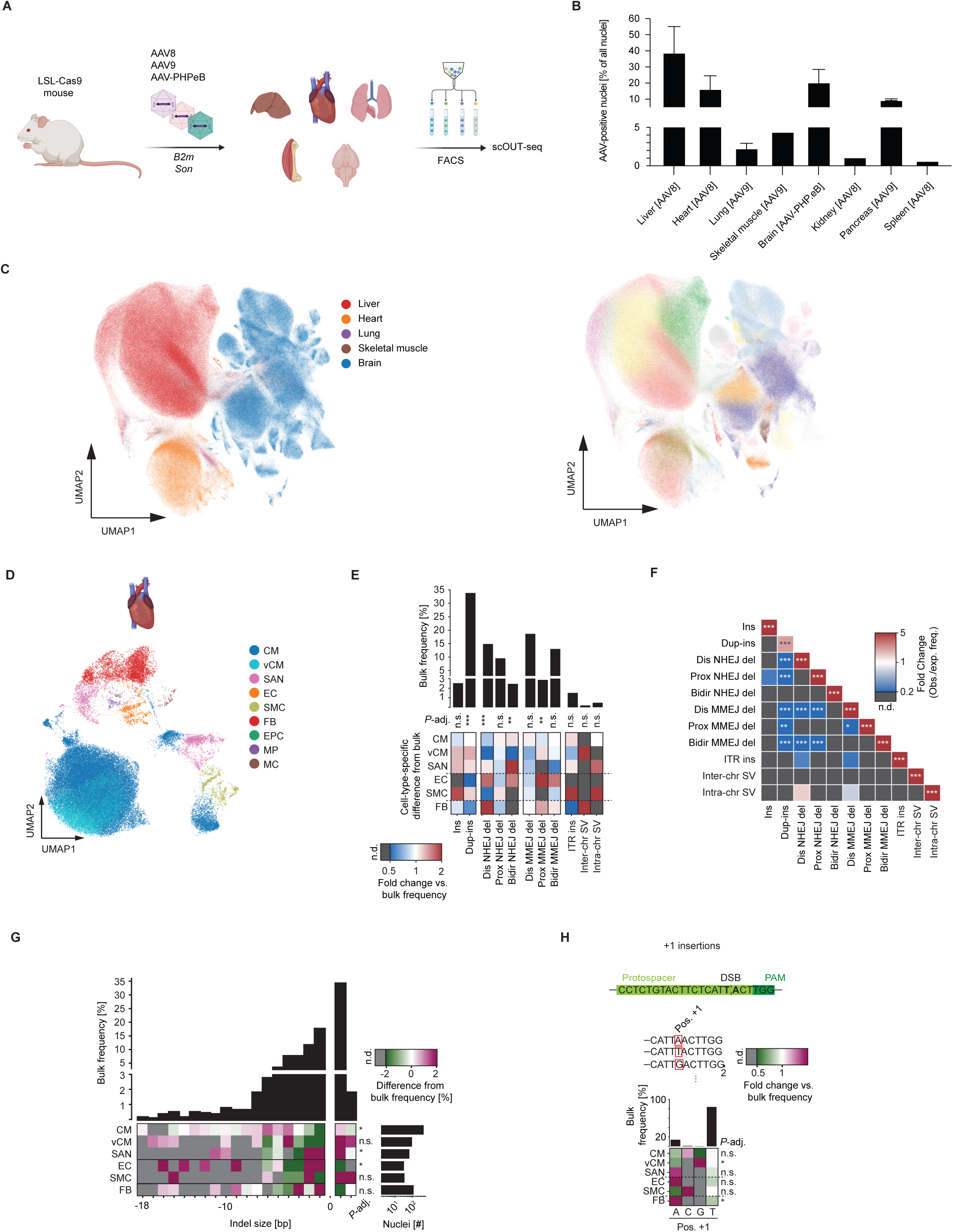
scOUT-seq across different tissues *in vivo*. (A) Schematic of mapping editing outcomes in different tissues following *in vivo* editing. AAVs were administered intravenously to the tail vein and organs were extracted 6 weeks post infection. (B) Bar graph depicting infection efficiencies across organs when using indicated AAV serotypes. Error bars represent the mean ± standard deviation (n = 3–5). (C) UMAP plots depicting tissue identities of 1,221,233 i*n vivo* edited nuclei (*left*, color coding for different tissues; *right*, color coding for different cell types). See **Figure S7A** for legend of cell types. (D) UMAP plot depicting 67,804 *in vivo* edited mouse heart nuclei colored by cell type, 1,582 of which were assigned with editing outcomes. CM, cardiomyocytes; vCM, ventral cardiomyocytes; SAN, sinoatrial node cells; EC, endothelial cells; SMC, smooth muscle cells; FB, fibroblasts; EPC, epicardial progenitor cells; MP, macrophages; MC, mural cells. (E) Bar graph showing bulk frequencies of editing outcome categories in mouse heart (*top*). Heatmap of cell-type-specific fold-changes relative to bulk frequencies. n.d., not detected. (*bottom*). FDR-adjusted p-values of G-tests determining difference of categories across cell lines are shown (**, *p*-adj. < 0.01; ***, *p*-adj. < 0.001; n.s., p-adj. > 0.05). (F) Correlation plot of fold-changes between observed and expected frequency of co-occurrence (assuming independence of alleles) of editing outcome categories in mouse heart. n.d., not detected. FDR-adjusted p-values of a χ² (chi-square) test determining difference between observed and expected frequencies across cell types are shown (*, *p*-adj. < 0.05; **, *p*-adj. < 0.01; ***, *p*-adj. < 0.001). (G) Histogram showing bulk frequencies of indel sizes in mouse heart (*top*). Heatmap of percentage-point deviations in indel-size frequencies from the bulk across cell types. n.d., not detected. (*bottom*). The number of cells per cell type is depicted in a bar graph. FDR-adjusted p-values of G-tests determining difference of indel profile to bulk are shown (*, *p*-adj. < 0.05; ***, *p*-adj. < 0.001; n.s., p-adj. > 0.05). (H) Insertion sequence composition of single base pair insertions in the mouse heart. The fold change compared to the bulk is plotted across cell types and insertion sequence positions. n.d., not detected. Bar graphs depict the bulk frequency of each base at each position. The sequence context of the *B2m* sgRNA is depicted as a reference. FDR-adjusted p-values of G-tests determining difference of insertion sequence composition to bulk are shown (*, *p*-adj. < 0.05; n.s., p-adj. > 0.05). See also **Figure S7**

To evaluate tissue-specific gene editing, we delivered guide RNAs against the 3′ untranslated region (UTR) of *B2m* to liver (three edited mice, one non-targeting control (NTC); AAV8), heart (two edited mice, one NTC; AAV8 and AAV9), lung (two edited mice, one NTC; AAV9) and skeletal muscle (two edited mice; AAV8 and AAV9). Because *B2m* is minimally expressed in the brain, the 3′ UTR of *Son* was targeted instead (five edited mice, two NTCs; AAV-PHP.eB). The AAV8, AAV9 and AAV-PHP.eB serotypes were chosen to maximize transduction efficiency in their target organs^46–49^. Infection efficiencies varied between organs and AAV serotypes (**Figure 5B).** Cargo delivery to liver, heart and brain was relatively efficient, while some organs like kidney or spleen were not infected well. The pancreas was infected at a reasonable efficiency, but RNA quality was relatively poor (**Data not shown**), potentially due to the enzymes produced by the organ. We therefore focused on the liver, heart, skeletal muscle, lung and brain, which all exhibited reasonably high infection rates and high-quality RNA preparations.

We dramatically increased cell throughput by integrating scOUT-seq with combinatorial indexing-based single-nucleus RNA-sequencing, enabling us to collect data on 1,221,233 total nuclei. After stringent filtering for unambiguous assignment of cell type and multi-allelic molecular genotypes, this yielded 349,725 editing outcomes within 182,898 transcriptionally assigned cells spanning 59 cell types (**Figure 5C, S7A**).

From skeletal muscle and lung, we recovered 15,581 and 9,479 cells, respectively, yielding 365 and 311 editing outcomes. Because these datasets are comparatively small, we focus subsequent analyses on tissues with higher coverage and present the skeletal muscle and lung results in the Supplement (**Figure S7B, C**).

### Editing outcomes in the mouse heart

We first investigated how editing outcomes vary between different cell types of the mouse heart. We profiled the transcriptomes of 67,804 nuclei spanning 9 cell types and assigned 3,012 molecular editing outcomes (**Figure 5D, S7D**). This number is relatively low compared to the number of analyzed nuclei potentially due to the low expression of *B2m* in heart cells (**Figure S7E**). Epicardial progenitor cells (EPC, 0 non-WT nuclei), macrophages (MP, 2 non-WT nuclei) and mural cells (MC, 4 non-WT nuclei) were not abundant enough to draw conclusions and were thus excluded from quantitative analysis.

Duplication-insertions were the most prevalent editing outcome in the heart (**Figure 5E**), consistent with observations in human HSPCs (**Figure 2B**). However, this is in contrast to human upper airway organoids, which preferentially exhibited distal NHEJ deletions (**Figure 2I**). Genome editing using AAV can lead to integration of viral inverted terminal repeats (ITRs) into the site of the DSB, sometimes accompanied by adjacent vector backbone or transgene sequences^50^. These ITR insertions were indeed detectable during *in vivo* editing, with variable frequency per cell type (**Figure 5E**).

When comparing the frequency of each editing outcome in individual heart cell types to the bulk, we found that SMCs were enriched for multiple types of insertions, especially ITR and non-templated insertions (**Figure 5E**). By contrast, ITR insertions were depleted from FBs and entirely missing from ECs. Several cell types exhibited a propensity for various types of deletions. ECs showed elevated levels of both PAM-proximal and bidirectional MMEJ deletions while SAN cells exhibited the highest levels of bidirectional NHEJ deletions and FBs were dominated by PAM-distal NHEJ deletions (**Figure 5E**). Intra-chromosomal SVs were most abundant in SMCs, while inter-chromosomal SVs were abundant in vCMs (**Figure 5E, S7F**). vCMs also showed a marked reduction in distal and bidirectional NHEJ deletions. CMs and ECs showed greater overall MMEJ deletion frequencies, which could be caused by their higher *Polq* expression than SMCs and SANs, which showed fewer MMEJ deletions (**Figure 5E, S7F, G**). Similar to results in human HSPCs and human UAOs, we found that homozygous outcomes occurred more frequently than expected (**Figure 5F**). However, unlike in other HSPCs and organoids, there were no significant exceptions to this pattern.

To investigate cell-type-specific differences in indel size profiles, we aggregated the molecular indel frequency distributions for each cardiac cell type and benchmarked them against the bulk profile. Relative to bulk, cardiomyocytes (CMs and vCMs) exhibited fewer 1–3-bp deletions and an enrichment of 4-bp deletions (**Figure 5G**). In contrast, FBs showed elevated 1-bp and 3-bp deletions alongside a reduction in 1-bp insertions (**Figure 5G**). Consistent with *ex vivo* results, most +1 insertions in the bulk population were T duplications (**Figure 5H**). When we stratified +1 insertions by cell type, we found that vCMs preferentially incorporated G, whereas FBs, ECs and SANs were enriched for A (**Figure 5H**). +2 insertions were observed too infrequently to draw robust conclusions, and +3 insertions were not detected at all (**Figure S7H, I**). These findings reveal cell-specific biases in DNA repair and editing outcome spectra in the heart, prompting us to ask whether similar patterns were present in other tissues.

### Editing outcomes in the mouse liver

We next investigated how editing outcomes vary between different cell types of the mouse liver. We edited the livers of 3 mice in the *B2m* 3’ UTR and assigned 10 distinct cell types from the transcriptomes of 467,743 nuclei, with 219,988 molecular editing outcomes (**Figure 6A, S8A**). As in other organs, we observed no effect of editing the 3’ UTR on the transcript levels of *B2m* (**Figure S8B**). We grouped the cell types of the liver into three broad classes: progenitors (HPC), parenchymal cells (Hep, PP-hep, Hyb-hep), and non-parenchymal cells (ChoC, SC, HepSC, LSEC, KC) (**Figure 6B**).

**Figure 6.**
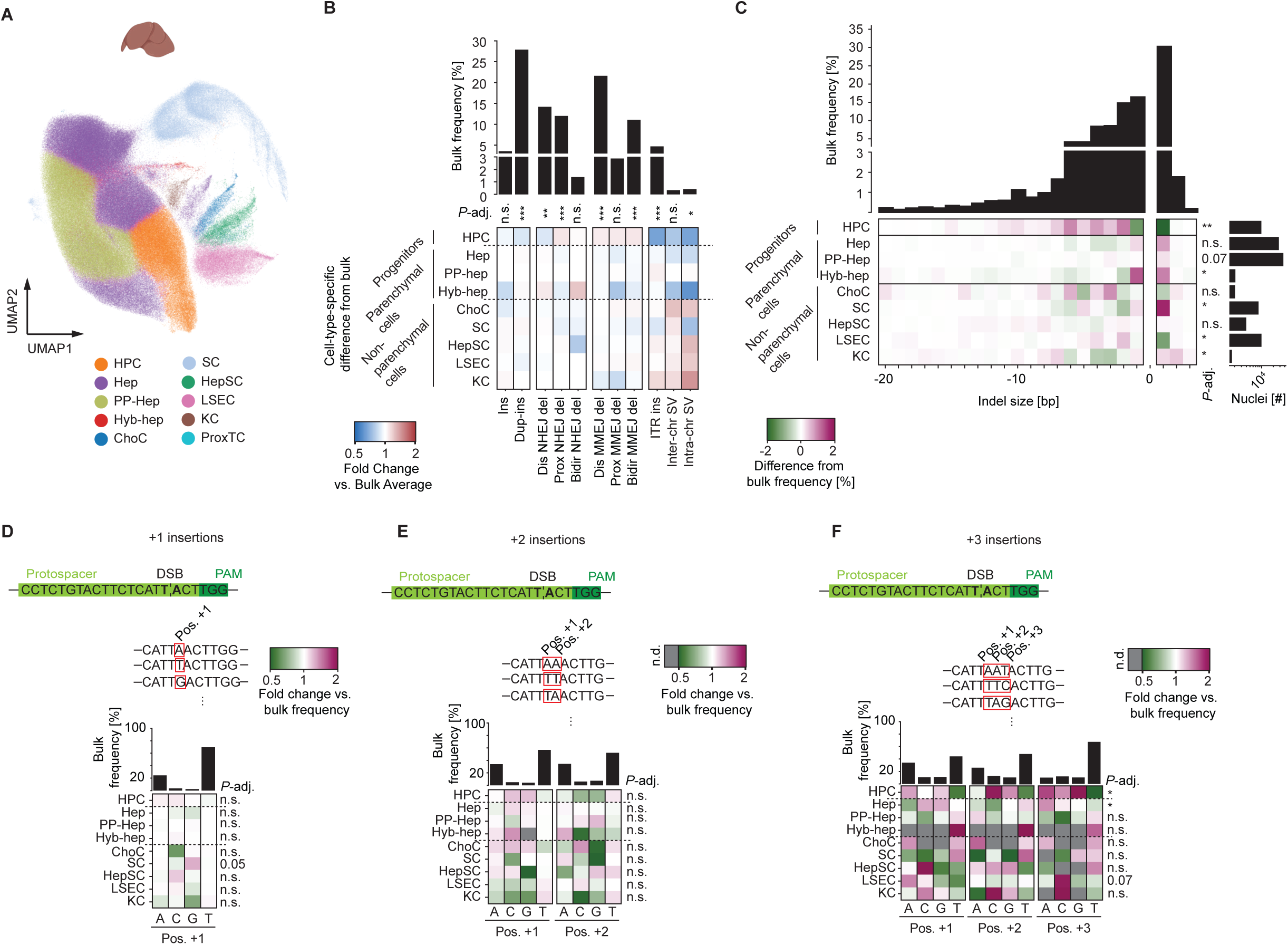
Mapping editing outcomes in mouse liver. (A) UMAP plot depicting 467,743 *in vivo* edited mouse liver nuclei colored by cell type, 106,817 of which were assigned with editing outcomes. HPC, hepatocyte progenitor cells; Hep, hepatocytes; PP-hep, periportal hepatocytes; Hyb-hep, hybrid hepatocytes; ChoC, cholangiocytes; SC, stromal cells; HepSC, hepatic stellate cells; LSEC, liver sinusoidal endothelial cells; KC, Kupffer cells; ProxTC, proximal tubular cells. (B) Bar graph of bulk frequencies of editing outcome categories in mouse liver (*top*). Heatmap of cell-type-specific fold-changes relative to bulk frequencies (*bottom*). FDR-adjusted p-values of G-tests determining difference of categories across cell lines are shown (**, *p*-adj. < 0.01; ***, *p*-adj. < 0.001***). (C) Histogram showing bulk frequencies of indel sizes in mouse liver (*top*). Heatmap of percentage-point deviations in indel-size frequencies from the bulk across cell types in mouse liver. The number of cells per cell type is depicted in a bar graph (*bottom*). FDR-adjusted p-values of G-tests determining difference of indel profile to bulk are shown (*, *p*-adj. < 0.05; ***, *p*-adj. < 0.001; n.s., p-adj. > 0.05). (D) Insertion sequence composition of +1 insertions in the mouse liver. The fold change compared to the bulk is plotted across cell types and insertion sequence positions. Bar graphs depict the bulk frequency of each base at each position (*top*). The sequence context of the *B2m* sgRNA is depicted as a reference. FDR-adjusted p-values of G-tests determining difference of insertion sequence composition to bulk are shown (*, *p*-adj. < 0.05; n.s., p-adj. > 0.05). (E) Insertion sequence composition of +2 insertions in the mouse liver. The fold change compared to the bulk is plotted across cell types and insertion sequence positions. n.d., not detected. Bar graphs depict the bulk frequency of each base at each position (*top*). FDR-adjusted p-values of G-tests determining difference of insertion sequence composition to bulk are shown (***, *p*-adj. < 0.001; n.s., p-adj. > 0.05). (F) Insertion sequence composition of +3 insertions in the mouse liver. The fold change compared to the bulk is plotted across cell types and insertion sequence positions. n.d., not detected. Bar graphs depict the bulk frequency of each base at each position (*top*). FDR-adjusted p-values of G-tests determining difference of insertion sequence composition to bulk are shown (*, *p*-adj. < 0.05; **, *p*-adj. < 0.01; ***, *p*-adj. < 0.001; n.s., p-adj. > 0.05). See also **Figure S8**.

Editing profiles were generally consistent within these classes, with some exceptions, but showed several clear distinctions between classes (**Figure 6B**). Notably, progenitors were slightly depleted for NHEJ-type alleles (insertions and NHEJ deletions), slightly enriched for MMEJ type alleles, and strong depleted for structural variants such as ITR insertions and translocations. This is in contrast to non-parenchymal cells, which generally enriched ITR insertions and both inter- and intra-chromosomal translocations. This could suggest a protective mechanism in hepatic progenitors and highlights the potential for unanticipated consequences of genome editing arising from difficult-to-detect structural variations in the non-parenchymal cells that comprise up to 30% of the liver.

To determine the differences in indel size profiles between cell types of the mouse liver, we aggregated indel size frequencies by cell type and compared them to the bulk frequencies. Within the non-parenchymal class, SCs and KCs were enriched for 1-bp insertions and displayed reduced frequencies of 1-4 bp deletions (**Figure 6C**). As opposed to multiple progenitor subtypes in the human HSPC sample, hepatic progenitors (HPC) were markedly depleted for 1-bp insertions and deletions. However, HPCs showed a concomitant enrichment of 2-8 bp deletions. These larger deletions form the basis of the shift away from classical NHEJ toward MMEJ in hepatic progenitors observed in the binned outcomes, despite their low *Polq* levels (**Figure 6B, S8C**). This shift potentially reflects distinct end resection propensity or expression of MMEJ-related repair factors other than *Polq* in hepatic progenitors compared to progenitor subtypes in the human HSPC samples.

As in other tissues, liver +1 insertions mostly duplicated the PAM-distal nucleotide next to the DSB (T), followed by duplication of the PAM-proximal nucleotide (A) (**Figure 6D**). Relative to bulk, cholangiocytes were depleted for C among +1 insertions, whereas stromal cells were enriched for G. For +2 insertions, the dinucleotide profile differences broadly fit into three main cell classes, with hepatic progenitors somewhat over-representing C/G at position 1 and under-representing C/G at position 2, while non-parenchymal cells under-represented C/G at both position 1 and position 2. (**Figure 6E**). +3 insertions in bulk displayed a more uniform base distribution (**Figure 6F**). Across non-parenchymal populations, G and T were consistently under-represented at position 1 and C was enriched at position 2, while hepatic progenitors were uniformly depleted of T across all three positions (**Figure 6F**). Together, these data indicate that the nucleotide composition of insertion sequences varies with both insertion length and cell identity between liver progenitors, parenchymal, and non-parenchymal cells, hinting at distinct insertion mechanisms.

### Editing outcomes in the mouse brain

Finally, we examined cell-type-specific differences between genome editing outcomes in the mouse brain. We performed *in vivo* editing of the 3’ UTR of the *Son* gene in 5 mice using the AAV-PHP.eB serotype. Overall, we profiled 523,029 brain nuclei spanning 19 cell types and assigned 126,049 molecular editing outcomes (**Figure 7A, S9A).** Consistent with reports in bulk tissues, NHEJ outcomes such as duplication-insertions and PAM-distal NHEJ deletions were the dominant edit types in the brain^51^ (**Figure 7B**).

**Figure 7.**
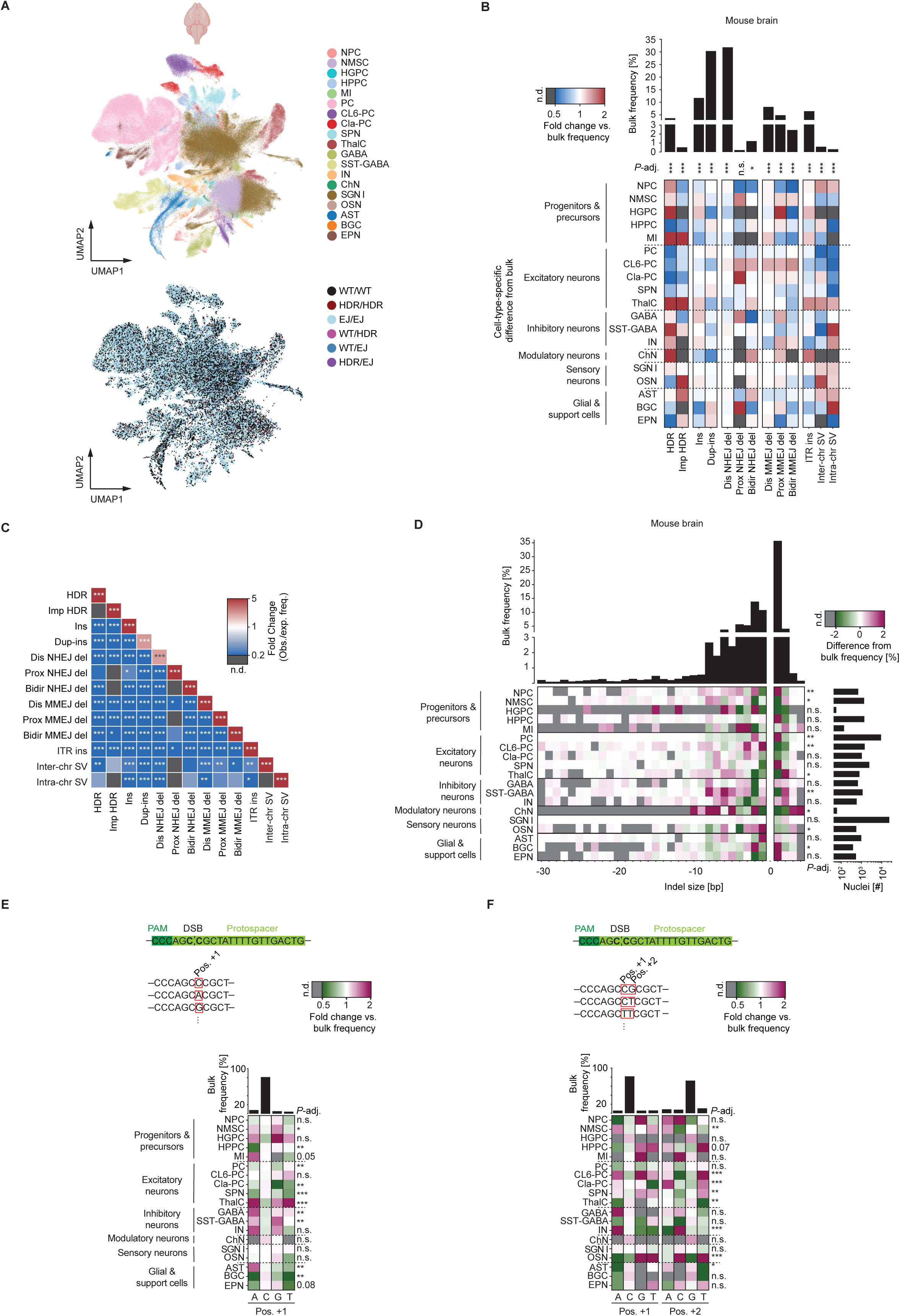
Mapping editing outcomes in mouse brain. (A) UMAP depicting cell identities of 523,029 *in vivo* edited mouse brain nuclei colored by cell type (*top*). UMAP of 64,936 mouse brain nuclei assigned with editing outcomes and colored by editing outcome category (*bottom*). NPC, neuronal progenitor cells; NMSC, non-myelinating stem cells; HGPC, hippocampal granule precursor cells; HPPC, hippocampal pyramidal precursor cells; MI, migrating interneurons; PC, pyramidal cells; CL6-PC, cortex layer 6 pyramidal cells; Cla-PC, claustrum pyramidal cells; SPN, subcortically projecting neurons; ThalC, thalamus cells; GABA, GABAergic neurons; SST-GABA, somatostatin GABAergic neurons; IN, inhibitory neurons; ChN, cholinergic neurons; SGN I, type I spiral ganglion neurons; OSN; olfactory sensory neurons; AST, astrocytes; BGC, Bergmann glial cells; EPN, ependymocytes; WT, wild type; HDR, homology-directed repair; EJ, end-joining. (B) Bar graph depicting bulk frequencies of editing outcome categories in mouse brain (*top*). Heatmap of cell-type-specific fold-changes relative to bulk frequencies. n.d., not detected. (*bottom*). FDR-adjusted p-values of G-tests determining difference of categories across cell lines are shown (*, p-adj. < 0.05; **; *p*-adj. < 0.01; ***; *p*-adj. < 0.001; n.s., p-adj. > 0.05). (C) Correlation plot of fold-changes between observed and expected frequency of co-occurrence (assuming independence of alleles) of editing outcome categories in mouse brain nuclei. n.d., not detected. FDR-adjusted p-values of a χ² (chi-square) test determining difference between observed and expected frequencies across cell types are shown (*, *p*-adj. < 0.05; **, p-adj < 0.01; ***, *p*-adj. < 0.001). (D) Histogram showing bulk frequencies of indel sizes in mouse brain (*top*). Heatmap of percentage-point deviations in indel-size frequencies from the bulk across cell types. n.d., not detected. (*bottom left*). The number of cells per cell type is depicted in a bar graph (*bottom right*). FDR-adjusted p-values of G-tests determining difference of indel profile to bulk are shown (*, *p*-adj. < 0.05; **, *p*-adj. < 0.01; ***, *p*-adj. < 0.001; n.s., p-adj. > 0.05). (E) Insertion sequence composition of +1 insertions in the mouse brain. The fold change compared to the bulk is plotted across cell types and insertion sequence positions. n.d., not detected. Bar graphs depict the bulk frequency of each base at each position (*top*). The sequence context of the *Son* sgRNA is depicted as a reference. FDR-adjusted p-values of G-tests determining difference of insertion sequence composition to bulk are shown (*, *p*-adj. < 0.05; **, *p*-adj. < 0.01; ***, *p*-adj. < 0.001; n.s., p-adj. > 0.05). (F) Insertion sequence composition of +2 insertions in the mouse brain. The fold change compared to the bulk is plotted across cell types and insertion sequence positions. n.d., not detected. Bar graphs depict the bulk frequency of each base at each position (*top*). FDR-adjusted p-values of G-tests determining difference of insertion sequence composition to bulk are shown (*, *p*-adj. < 0.05; **, *p*-adj. < 0.01; ***, *p*-adj. < 0.001; n.s., p-adj. > 0.05). See also Figure S9.

To determine whether editing outcome patterns correlated with cell function, we grouped brain cell types into six broad functional classes: progenitors/precursors (Neural progenitor cells, non-myelinating stem cells, hippocampal granule precursor cells, hippocampal pyramidal precursor cells, migrating interneurons), excitatory neurons (pyramidal cells, cortex layer 6 pyramidal cells, claustrum pyramidal cells, subcortically projecting neurons, thalamus cells), inhibitory neurons (GABAergic neurons, somatostatin-GABAergic neurons, inhibitory neurons), modulatory neurons (Cholinergic neurons), sensory neurons (Type I spiral ganglion neurons, olfactory sensory neurons), and glial/support cells (Astrocytes, Bergmann glial cells, ependymocytes) (**Figure 7B**). We found that several binned outcomes showed consistent patterns within classes.

Target-anchored inter-chromosomal SVs were preferentially detected in sensory neurons, whereas intra-chromosomal SVs were enriched in sensory neurons, inhibitory neurons, and glial/support cells. Conversely, intra-chromosomal SVs were depleted in excitatory neurons (**Figure 7B**). We found HDR alleles enriched in progenitor/precursor cells, consistent with the mitotic potential of these cell types. Most post-mitotic neuronal cell types under-represented HDR alleles. But we were surprised to find HDR alleles over-represented in post-mitotic inhibitory and modulatory neuron populations (**Figure 7B**). For example, allele tables derived from Cla-PC and SST-GABA cell reads revealed a marked increase in the proportion of clear HDR reads in SST-GABA cells, while the proportion of unedited reads remained nearly constant (**Figure S9B**). Post-mitotic sensory neurons and mitotic glial/support cells under-represented perfect HDR but favored imperfect HDR events that represented partial conversion of the template.

Some cell types diverged substantially from the rest of their functional class. Unlike most progenitor/precursors, HPPCs were markedly depleted for HDR alleles (**Figure 7B**). Unlike other excitatory neurons, CL6-PC cells exhibited elevated levels of all types of NHEJ and MMEJ deletions, whereas ThalC cells were enriched for both inter- and intra-chromosomal SVs as well as HDR and imperfect HDR. Notably, cell types with increased ITR insertions typically also showed higher frequencies of perfect HDR (e.g., HGPC, MI, ThalC, and ChN). In the brain as in the liver, *Polq* expression level and prevalence showed surprisingly poor correlation with MMEJ deletion levels **(Figure S9C**). Collectively, these data show that while certain editing patterns are shared among functionally related cell types, broadly descriptive cell function alone does not fully account for the diversity of editing outcomes.

To understand which editing outcomes co-occur with each other in the same cell, we calculated the expected co-occurrence frequencies assuming independence of outcomes and compared it with the observed co-occurrence frequencies in our brain sample. Despite targeting a distinct locus (*Son* 3’ UTR) than the mouse heart (*B2m* 3′ UTR) and in human HSPCs and UAOs, the mouse brain also displayed a significantly higher frequency of homozygous edits than expected (**Figure 7C**). These outcomes were generally identical in both binned type and molecular sequence, as uniquely assignable from a combination of cell barcode and UMI. As in other cell types, this suggests that cells are either predisposed to similar molecular outcomes or can copy the repair outcome from an already-repaired allele to an as-yet unrepaired allele.

Turning to molecular indel profiles of the brain cell classes, we found that the indel size distributions of most cell types deviated from the bulk average (**Figure 7D**). Indel spectra were relatively coherent within inhibitory neurons and glial/support cells. Inhibitory neurons were depleted for 2 bp deletions but enriched for deletions of 3-9 bp and for 2 bp insertions. Conversely, glial/support cells were enriched for 1-bp insertions. Modulatory neurons also markedly diverged from the bulk indel profile, with substantial enrichment for 5-10 bp deletions, 2 bp deletions and 3-4 bp insertions, alongside depletion of 1 bp deletions and 1-2 bp insertions.

However, within broader functional classes, several cell types deviated from class trends (**Figure 7D**). Among progenitors/precursors, NPCs preferentially accumulated intermediate-length deletions (4-9 bp) similar to INs, and MIs were enriched for 3-4 bp deletions. Among excitatory neurons, PCs favored 1 bp insertions and 2 bp deletions while being comparatively depleted of 3-8 bp deletions. Within sensory neurons, OSNs were enriched for 1-3 bp deletions, whereas SGN I cells resembled the bulk profile, likely reflecting their high abundance. These data suggest that while functionally similar cell types share certain molecular editing outcomes, several features are cell-type-specific.

We compared the composition of +1, +2, and +3 insertion sequences across cell classes and subtypes. Consistent with the *ex vivo* samples, the vast majority of +1 insertions templated the PAM-distal nucleotide at the cut site, and this also held true for the first position of +2 and +3 insertions (**Figure 7E-F, S9D**). However, non-templated +1 insertions exhibited substantial cell type stereotyping. Multiple inhibitory neuron subtypes exhibited more A and G and fewer C insertions than the bulk average. For +2 insertions, most excitatory neuron types showed an increased frequency of T and a concomitant depletion of C at the second position relative to bulk. By contrast, +2 insertions in IN and GABA cells were enriched for A at the first position and C at the second position. Plus 3 insertion sequence composition appeared to vary in certain cell types (NMSCs, HPPCs, ChNs, BGCs), but small group sizes limited power and preclude firm conclusions (**Figure S9D, E**). Together, these data reveal cell-type- and functional-group-specific biases in insertion sequence composition, consistent with selective engagement of distinct DNA repair pathways.

Collectively, our data establish scOUT-seq as a robust platform to investigate single cell genome editing outcomes in diverse cell types, editing modalities, and organisms. Our observations across multiple *ex vivo* and *in vivo* settings highlight the impact of cell type on Cas9-mediated genome editing and provides large-scale initial maps of preferential and disfavored editing outcomes in numerous contexts.

## Discussion

Here, we describe scOUT-seq, an experimental and computational approach to map molecular genome editing outcomes, cell type, and state at single cell and single nucleotide resolution. We applied scOUT-seq to investigate editing profiles in heterogeneous cell mixtures of both human and mouse cells, including *ex vivo* and *in vivo* edited samples processed with a variety of single cell preparation technologies. We found that in most cases the bulk editing profile reflects only the most prevalent cell type within a complex mixture, and that cell type, state, and target sequence context shape molecular editing outcomes. These data are available in an easily browsable form at [XYZ].

By measuring repair at a Cas9 target site located within a locus highly expressed across all cells, we normalized sequence-context effects and reduced, though potentially not eliminated, chromatin and 3D-architectural confounders. In several cases, less differentiated cells showed more small deletions (<7 bp) and fewer +1 insertions, whereas more differentiated cell types accumulated large deletions (>7 bp) consistent with a shift from classical NHEJ towards resection-dependent MMEJ. Mechanistically, this bias could reflect increased chromatin compaction in differentiated cells that restricts access of NHEJ factors (Ku70/Ku80) and/or altered levels of resection-associated proteins including CtIP and the MRN complex^52^. Indeed, chromatin context and higher-order genome architecture modulate repair factor access, γH2AX establishment and DDR focus assembly^13,53,54^. We found that POLQ expression tracked MMEJ outcomes well in some tissues but poorly in others, implicating additional regulators. These results align with systematic perturbation maps that resolved two genetically distinct mechanisms of MMEJ downstream of MRE11-mediated end processing: a POLQ-dependent mode promoted by RAD17 and the 9-1-1 complex and a POLQ-independent route^21^. The shallow nature of polyA-based single cell RNA sequencing limits the ability of scOUT-seq itself to comprehensively read out the levels of the hundreds of known DDR factors, which are anyway subject to extensive post-transcriptional and post-translational regulation. The multi-faceted links between differential repair factor activity and differential editing outcomes remain to be elucidated.

Studies in homogeneous cell source show that inhibition of DNA-PKcs suppresses classical NHEJ while increasing HDR, translocations and large deletions^41,55^. Consistent with this, we observed that target-anchored intra-chromosomal structural variants were generally induced by DNA-PKcs inhibition across multiple cell types. Detecting structural variants including large deletions, inversions, and translocations remains difficult with both short- and long-read sequencing approaches because breakpoints are often unpredictable and can occur megabases apart or on different chromosomes. Using scOUT-seq, we found that the extent to which DNA-PKcs inhibition altered the frequency of SVs was cell type specific. For example, intrachromosomal SVs rose in CD34+ lymphoid-primed HSCs and organoid-derived hillock cells upon DNA-PKcs inhibition, whereas they declined in CD34+ myeloid-primed HSCs and organoid-derived ionocytes. This implies that the NHEJ backup pathways that can lead to SVs are differentially active in various cell types.

Hillock cells are considered more injury resistant to both physical and chemical damage and serve as a plastic reservoir that aids tissue repair^56^. This intrinsic resilience may also enable these cells to better tolerate large genomic perturbations. Notably, increases in intra-chromosomal SVs did not necessarily coincide with increases in inter-chromosomal SVs. For example, DNA-PKcs inhibition reduced target-anchored inter-chromosomal structural variants in human HSPCs. This could be due to increased reliance on canonical NHEJ, which promotes translocations in human cells^44^.

We were surprised to detect clear HDR signatures in the brain. These were apparent not only in neural progenitor populations, but also in several differentiated inhibitory neuronal subtypes, thalamic, and cholinergic neurons. Observing HDR outcomes in mature neurons is intriguing as it raises the possibility that this precise DDR pathway could be used to address certain brain disorders. However, it is not yet clear whether these outcomes reflect HDR in rare dividing progenitors that differentiate to certain neuronal subtypes, a subset of neurons capable of limited cell-cycle re-entry and re-quiescence, or yet-uncharacterized HDR-like processes in post-mitotic cells^57–59^. Reports have suggested that differentiated neurons may transiently re-enter the cell cycle in response to genotoxic stress in order to access repair pathways typically reserved for proliferating cells^60^. This parallels transient cycling to enable HDR in engrafting LT-HSCs from the bone marrow^31^. Such a mechanism could have implications for both genome stability and neurodegeneration, as inappropriate or incomplete cell-cycle re-entry is linked to neuronal dysfunction and cell death^60^. Better understanding of the origin of HDR in neuronal subpopulations would help understand how the nervous system balances genome maintenance with functional stability, and to guide the design of future editing therapies targeting neural tissues.

The simultaneous capture of editing outcome and cell identity make scOUT-seq a powerful tool to unravel the cell-type-specificity of genome editing tools both *in vivo* and *ex vivo*. Here we focused on large-scale maps to reveal the extent of differential genome editing with correlative relationships between cell types and outcomes. Machine-learning approaches trained in immortalized cells can predict editing outcomes and direct guide RNA choice towards those with desired sequence outcomes^17,61–63^. Our data indicate that these sequence outcomes may broadly apply, but their prevalence in a cell type of interest could be markedly higher or lower than that predicted by models trained on bulk outcomes in immortalized cells. This would lead to missed opportunities, for example choosing an editing reagent to yield a therapeutic outcome when a different reagent would be more effective in the relevant cell type. A great deal more data spanning many sequence contexts and for each cell type of interest would be needed to enable automated prediction of cell stereotyped differential editing outcomes. However, large-scale perturbation approaches could determine causation for differential editing outcomes and potentially allow one to modulate general predictions based on known expression patterns for target cell types.

The repair of DNA damage caused by different kinds of editors is complex and remains difficult to predict. The large-scale maps provided here will hopefully help design genome editing interventions for target tissues and cell types, and continued use of scOUT-seq would further expand these resources for the research and preclinical communities.

## Methods

### Cell culture

Murine fibroblast cells (NIH3T3) were purchased from the German Collection of Microorganisms and Cell Cultures GmbH and tested for mycoplasma. They were cultured in DMEM, supplemented with 10% FBS and 100 µg ml−1 penicillin–streptomycin.

Murine myoblast cells (C2C12) were obtained from the Gehart lab and tested for mycoplasma. They were cultured in DMEM, supplemented with 10% FBS and 100 µg ml−1 penicillin–streptomycin.

Murine neuroblast cells (Neuro-2a) were purchased from the German Collection of Microorganisms and Cell Cultures GmbH and cultured in EMEM, supplemented with 10% FBS and 100 µg ml−1 penicillin–streptomycin.

Human G-CSF-mobilized CD34^+^ HSPCs from adult healthy donors were purchased from the Fred Hutchinson Cancer Center and cultured in StemSpan SFEM II media (STEMCELL Technologies, 09655) supplemented with StemSpan CC110 (1×) (STEMCELL Technologies, 02697) and 100 µg ml−1 penicillin–streptomycin.

The human primary nasal epithelial samples were collected from healthy volunteers, in accordance with ethical guidelines of the ETH Zürich Ethics Commission (EK-2024-N-171-A) and the Cantonal Ethics Committee Zürich (Req-2O24-OO558). The collected tissue was digested into single cells using TrypLE (Thermo Fisher Scientific, 12605010) and seeded in 90% BME drops (Biotechne, 3533-005-02) with mLung+ medium supplemented with Y-27632 (Tocris, 1254). Cultures were maintained with medium changes every 2–3 days until organoid structures formed. During routine culture, human upper airway organoids were seeded in six-well plates in 15 μl of BME droplets and polymerized for 30 min at 37 °C. After BME solidification, organoid-specific culture medium (**Table S1**) was added, and the organoids were cultured at 37 °C until further use. For each passage (every 4–6 d), the organoids were dissociated mechanically, washed to remove remaining BME and seeded at half the density in fresh BME.

### Murine LSK isolation and culture

Bone marrow was isolated from the hind limbs (femurs and tibias) of 5 Cas9-EGFP C57BL/6 female mice by trimming bone ends and flushing the marrow with PBS. Red blood cells were lysed with ACK lysis buffer (ThermoFisherScientific, A1049201) and Lin (-) cells were extracted from the mixture using MACS depletion of lineage cells (Miltenyi Biotec, 130-110-470). The Sca-1⁺/c-Kit⁻ phenotype was sorted by FACS using PE-Cy7-Sca (Clone D7, ThermoFisher, 25-5981-82) and APC-c-kit (Clone ACK2, Biolegend, 135107) for staining. Upon isolation, cells were cultured in 96 well plates in IMDM with 20% BIT, 1% P/S, 100 ng/ml TPO and 100 ng/ml SCF.

### Cas9 RNP generation

For the cell line mixing experiment, 120 pmol of Gpi1 sgRNA and 120 pmol of Pgk1 sgRNA were mixed with 200 pmol in-house produced SpCas9–NLS and 100 pmol of a Gpi1 single-stranded HDR template and 100 pmol of a Pgk1 single-stranded HDR template. The sgRNAs were produced by in vitro transcription (IVT) as described here: https://doi.org/10.17504/protocols.io.n2bvjyp5vk5w/v17. For IVT, overlapping oligomers containing a T7 promoter, the desired protospacer and gRNA scaffold were amplified using Phusion polymerase (New England Biolabs (NEB), M0530L). The unpurified DNA product was then in vitro transcribed using an NEB HiScribe T7 High Yield RNA Synthesis Kit (NEB, E2040L) by incubating at 37 °C for 16 h. The following day, the resulting RNA was treated first with DNase I followed by dephosphorylation by a heat-labile version of calf intestinal alkaline phosphatase (NEB, M0525L), purified with an miRNeasy kit (Qiagen, 217084), concentration measured by a NanoDrop 8000 spectrophotometer and frozen at −80 °C. Single-stranded HDR templates (ssODNs) were ordered from IDT (Ultramer IDT Oligos).

For editing human HSPCs and upper airway organoids in Nucleocuvettes, 600 pmol of *GAPDH* sgRNA and 500 pmol of ssODN HDR template were mixed with 500 pmol in-house produced SpCas9-NLS. The *GAPDH* sgRNA was ordered from Synthego (CRISPRevolution sgRNA EZ Kit). The ssODN HDR template contained a 12-nucleotide substitution sequence around the cut site as well as a 2-nucleotide substitution to mutate the PAM, and a total length of 162 nucleotides.

### Sentinel sgRNA screening

To identify suitable sentinel sgRNAs in mouse cells, we first designed candidates targeting the 3′ UTRs of genes with high expression across diverse tissues and cell types. Candidate sgRNAs were then screened in fibroblasts and neuroblasts for high frequencies of end-joining repair outcomes using pooled amplicon-NGS analyzed with CRISPResso2^28^, prioritizing guides exhibiting a diverse spectrum of editing events. We subsequently tested selected sgRNAs in combination with ssODN HDR templates to quantify HDR rates in fibroblasts, neuroblasts, and myoblasts. We selected sgRNAs that achieved ≥10% HDR efficiency and ≥40% end-joining efficiency in at least one cell type.

In human cells, we similarly designed sgRNAs targeting the 3′ UTRs of highly expressed genes across diverse tissues and cell types. Initial screening in human CD34^+^ HSPCs assessed editing efficiency and outcome diversity using Sanger sequencing analyzed by TIDE^46^. To quantify HDR rates, we subsequently tested selected sgRNAs with ssODN HDR templates in CD34^+^ HSPCs and quantified outcome profiles using amplicon-NGS and CRISPResso2^28^. We again selected sgRNAs that achieved ≥10% HDR efficiency and ≥40% end-joining efficiency. A follow-up screening round in in human CD34^+^ HSPCs using newly designed sgRNAs and amplicon-NGS/CRISPResso2 analysis was performed to expand the pool of validated candidates for broader applicability across tissues and cell types^28^.

### Cell line mixing experiment

Murine fibroblast (NIH3T3), myoblast (C2C12) and neuroblast (Neuro-2a) cell lines were thawed and cultured in their respective media. Each cell line was then trypsinized and edited separately. 2×10^5^ NIH3T3 cells were resuspended in Lonza SG Nucleofection Buffer, 2×10^5^ C2C12 cells were resuspended in Lonza SE Nucleofection Buffer and 2×10^5^ Neuro-2a cells were resuspended in Lonza SF Nucleofection Buffer. The *Gpi1*/*Pgk1* Cas9-RNP/HDR template mix was then added to each cell line. The HDR template contained a constant 12 nucleotide sequence at the cut site between the homology arms to uniquely tag HDR events. The cell lines were electroporated in 20 µl strips (2 × 10^5^ cells per well). Immediately after electroporation, 80 µl of culture medium was added to each well and cells were allowed to recover for 5 min at 37 °C before being transferred to 24-well plates.

72 hours after electroporation, genomic DNA was extracted from the bulk population of each cell line using QuickExtractTM DNA Extraction Solution (Lucigen, QE0905T). The regions targeted by the *Gpi1*/*Pgk1* sgRNAs were then amplified using AmpliTaq Gold™ DNA-Polymerase (FisherScientific, 10428444) and primers including Illumina adapter sequences (25 PCR cycles). Indexing PCRs were then performed by amplifying (5 cycles) 100 ng of generated PCR products using Q5 High-Fidelity 2× PCR Master Mix (NEB, M0492L) and primers including i7 / i5 Illumina indexes. PCR products were purified using 1X SPRIselect beads (Beckman Coulter, B23318) after each PCR and quantified using Quant-iT dsDNA Assay Kits (Thermo Fisher Scientific, Q33120). Amplicon libraries were sequenced using 2 × 150 paired-end mode (Illumina) with a target average read count per amplicon of 200,000 reads. Illumina reads were demultiplexed and analyzed using the described scOUT-seq pipeline modified for bulk samples.

The remaining cells were mixed retaining an equal ratio between the cell lines, and 16000 cells of the mixture were analyzed using Chromium Controller (Firmware version 4.0) and Chromium Next GEM Single Cell 3′ Reagent Kits version 3.1 (10x Genomics) according to the manufacturer’s specifications. scRNA-seq Libraries were sequenced by the FGCZ.

scRNA-seq data were processed using Cell Ranger 6.0 with default parameters. Reads were aligned to mouse reference mm10. The Seurat (version 5.1.0) package was used to filter out cell multiplets, empty droplets and unhealthy cells by setting thresholds for number of reads per cell (200 < x < 6500) and percentage of mitochondrial reads per cell (<10%). Cell cycle phases scores were calculated and regressed out to mitigate the effects of cell cycle heterogeneity within cell lines. A K-nearest neighbor graph was constructed based on the euclidean distance in principal component space using the 2000 most variable genes and the first 12 principal components. To cluster the cells, the Louvain algorithm was applied to iteratively group cells together in order to optimize the standard modularity function. A resolution parameter of 0.5 was used. Then, Uniform Manifold Approximation and Projection (UMAP) was used to visualize the different clusters in low-dimensional space. Marker genes were identified for each cluster using differential gene expression analysis. These marker genes were used to assign cell line identity to each cluster. Allele distribution was then compared between bulk amplicon-NGS data per cell line and single cell assignments. The karyotyping of NIH-3T3 shows 4 copies of the *Gpi1* locus and 3 copies of the *Pgk1 locus* per cell^65^. In tetraploid C2C12, 4 copies of *Gpi1* and *Pgk1* are present^66^. Neuro-2a cells show an unstable karyotype with most cells containing 5 copies of *Gpi1* and *Pgk1*^67^. These copy numbers were assumed when assigning editing outcomes to the cells. To assign simplified editing outcomes for visualisation in the UMAP, threshold values for WT (Unedited), EJ (End-joining) and HDR percentage per cell were used: If more than 60% of alleles are WT, HDR or EJ, the cell is assigned as WT/WT, HDR/HDR or EJ/EJ, respectively. If 40-60% are WT and 40-60% are HDR, the cell is assigned as WT/HDR, and analogously WT/EJ and HDR/EJ are assigned.

### scOUT-seq 10X-based library generation

The standard 10x Genomics Chromium 3′ v.3 libraries were generated according to the manufacturer’s recommendations until recovery of first strand cDNA. The targeted *Gpi1* and *Pgk1* loci were amplified from 250 ng of cDNA using reverse gene-specific primers in combination with a generic forward sample index (SI)-PCR primer. The gene-specific reverse primers contain a partial Illumina read 2 handle. The SI-PCR oligonucleotide anneals to the partial Illumina read 1 sequence at the 3′ end of the molecule which preserves the cell barcode and UMI. The number of cycles used in PCR1 was determined experimentally and was dependent on the level of expression of the targeted gene (14 cycles for *Gpi1*, 11 cycles for *Pgk1*). After PCR1 unincorporated primers were removed using SPRI bead purification. A subsequent second PCR was performed with a sample indexing primer (RPI-X) and a universal forward PCR primer to retain cell barcode and UMI information and index the samples. scOUT libraries were quantified using Quant-iT dsDAssay Kits (Thermo Fisher Scientific, Q33120) and their size distribution/molarity was assessed using D1000 ScreenTape on an Agilent TapeStation. scOUT libraries were sequenced on different Illumina systems together with their parent 10x libraries to ensure sufficient sequence diversity. The following cycle settings were used for the mixing of cell lines experiment: 28 cycles for read 1, 91 cycles for read 2, and 8 cycles for the sample index.

For GAPDH targeting in human HSPCs and upper airway organoids, 50 ng of cDNA was used to amplify the targeted locus. PCR1 was performed with 17 cycles, the indexing PCR was performed with 6 cycles, and the following cycle settings were used to sequence the scOUT libraries. For human HSPCs: 28 cycles for read 1, 151 cycles for read 2, and 8 cycles for the sample index. For human upper airway organoids: 28 cycles for read 1, 99 cycles for read 2, and 8 cycles for the sample index.

### scOUT data analysis

During analysis, the scOUT library reads were used to generate a whitelist of the most likely true cell barcodes. The whitelist was generated using umi_tools^27^. Barcodes were parsed from the R1 reads according to the specified extraction method and barcode pattern. The frequency of each distinct barcode across the dataset was then counted. To distinguish true cell-associated barcodes from background noise, the distribution of barcode counts was analyzed using the density-based knee method. This approach fits the barcode rank–frequency distribution and identifies the inflection (“knee”) that separates abundant barcodes (putative real cells) from low-frequency barcodes (sequencing errors or empty droplets). The expected number of cells was provided as a prior to guide the cutoff, ensuring that the number of barcodes retained was consistent with the known complexity of the library. Nearly all reads (up to 999,999,999) were included in this estimation to maximize accuracy. Barcodes above the identified threshold were written to a whitelist file which was subsequently used to restrict downstream analyses to high-confidence cell barcodes. Barcodes/UMIs were then extracted and assigned to reads, and reads were discarded or corrected based on the whitelist. Reads were then aligned to the target reference using the CRISPResso algorithm (v2.3.1). For the *GAPDH* targeting samples, reads from known off target amplification events were filtered out using a blacklist of sequences. The aligned reads were then parsed and reads that had an imperfect priming sequence were discarded as off-target amplifications. Reads were classified by repair outcome and filtered according to their alignment scores to remove misaligned (misprimed) reads. Potentially misaligned reads were then further filtered. Misaligned reads in which the first mismatch occurred in the first 30% of the read were labelled as off-target amplifications. The other misaligned reads - reads that had an alignment score that was below what would be expected with their set of edits - were then split up into the portion aligning to the target and the translocation read portion. The translocated section of those reads was then aligned to the genome (hg38 for human samples, mm10 for mouse samples) using samtools (v1.13) and quantified by location and cell barcode/UMI combination^68^. Repair outcomes were assigned by determining repair outcomes for each UMI and then integrating those UMI-repair outcomes at the cell barcode level.

scOUT reads were categorized by the location and number of insertions, deletions and substitutions caused by the DSB. Furthermore, they were also divided into the following repair outcome categories. Reads containing a deletion without microhomology between the deleted sequence and the sequence surrounding the deletion were classified as non-homologous end joining (NHEJ-) deletions. If a microhomology of 2-25 nucleotides exists between the deleted sequence and the sequence surrounding the deletion, reads were classified as microhomology-mediated end joining (MMEJ-) deletions. Reads with both inserted and deleted bases were classified as “Deletions + Insertion”. Reads with inserted bases were generally classified as “Insertion”. If the inserted bases are duplicating the bases surrounding the insertion, the read was classified as duplication insertion. If the HDR template was correctly integrated, the read was classified as HDR. If the integration was imperfect, the read was classified as imperfect HDR. Reads that partially aligned to the targeted locus and a locus on another chromosome were classified as target-anchored inter-chromosomal structural variants, whereas reads that partially aligned to the targeted locus and the same chromosome were classified as target-anchored intra-chromosomal structural variants.

Cell type identity in combination with the distribution of read counts and UMI counts of the repair outcomes were used to determine the number of alleles to be assigned. Alleles were then assigned proportionally to the relative number of reads of each repair outcome on a per cell basis.

### Editing of human HSPCs

Human CD34^+^ HSPCs were thawed and cultured at a concentration of 0.7 × 10^6^ cells per milliliter for 24 hours before electroporation. Cells were resuspended in Lonza P3 Primary Cell Nucleofection solution and GAPDH RNP/HDR template mix was added to the cells. The cells were then electroporated in 100-µl single Nucleocuvettes (1 × 10^6^ cells per Nucleocuvette). Immediately after electroporation, 80 µl of culture medium was added to each well, and cells were allowed to recover for 5 min at 37 °C before being transferred to 6-well plates containing medium (final concentration of 1 × 10^6^ cells per milliliter) and 1 µM AZD7648 where indicated. Twenty-four hours after electroporation, cells were diluted by the addition of 1 ml of culture medium. 48 hours after electroporation, cells were sorted into a CD34^+^CD38^+^ progenitor population and CD34^+^CD38^-^ engraftment-enriched population to enrich for rarer cell types such as HSCs^30^. Gene expression and scOUT libraries were generated from the sorted populations as described above.

### Editing of human upper airway organoids

Human upper airway organoids were extracted from BME by pre-treatment with 2 U ml−1 Dispase II (Gibco, 17105041) for 30 min at 37 °C, followed by mechanical disruption and digestion with TrypLE (Gibco, 12604021) for 30 min at 37 °C. After digestion, cells were washed and strained through a 70-µm strainer and counted. In total, 1 × 10^6^ cells were then washed with PBS and resuspended in Lonza SE nucleofection buffer. Cas9 RNP/HDR template mix targeting *GAPDH* was added to the cells and the cells were electroporated. For cells to recover after the electroporation, complete warm culture medium was added to the cuvettes, and cells were incubated for 15 min at room temperature. After that, cells were transferred from the cuvettes to microcentrifuge tubes and pelleted (300g for 3 min). Afterwards, the cell pellet was resuspended in cold BME and plated as drops containing 25–100 × 10^5^ cells per drop. The cells were then cultured in complete culture medium containing 10 µM Y-27632 (TOCRIS, 1254) with or without AZD7648. After 67 hours, cells were extracted from BME by mechanical disruption and TrypLE digestion and scOUT-/scRNA-seq libraries were generated as described above.

### scRNA-seq data analysis and cell type assignment of human HSPCs and upper airway organoids

scRNA-seq data were processed using Cell Ranger 6.0 with default parameters. Reads were aligned to the human reference hg38. Seurat (version 5.1.0) was used to filter out cell multiplets, empty droplets and unhealthy cells by setting thresholds for number of reads per cell (1000 < x < 7500) and percentage of mitochondrial reads per cell (<20%). A K-nearest neighbor graph was constructed based on the euclidean distance in principal component space using the 2000 most variable genes and the first 30 principal components. To cluster the cells, the Louvain algorithm was applied to iteratively group cells together in order to optimize the standard modularity function. A resolution parameter of 1.2 was used. Then, Uniform Manifold Approximation and Projection (UMAP) was used to visualize the different clusters in low-dimensional space.

Each cell was scored for every possible cell type based on positive marker gene enrichment and negative marker absence using the CellMarker 2.0 database^69^. Cell type scores were then summed up across all cells in a cluster, and the top 3 cell types were evaluated manually before assigning a cell type to the cluster (**Table S2, 3**).

### scRNA-seq data analysis and cell type assignment in mouse LSK cells

10x scRNA-seq data were processed using Cell Ranger 6.0 with default parameters. Reads were aligned to the mouse reference mm10. Seurat (v5.1.0) was used to filter out cell multiplets, empty droplets and unhealthy cells by setting thresholds for number of reads per cell (1000 < x < 7500) and percentage of mitochondrial reads per cell (<20%). A K-nearest neighbor graph was constructed based on the euclidean distance in principal component space using the 2000 most variable genes and the first 30 principal components. To cluster the cells, the Louvain algorithm was applied to iteratively group cells together in order to optimize the standard modularity function. A resolution parameter of 1 was used. Then, Uniform Manifold Approximation and Projection (UMAP) was used to visualize the different clusters in low-dimensional space.

Evercode^TM^ whole transcriptome data was processed using the ParseBiosciences Pipeline (v1.3.1) with v2 chemistry and aligning against the mm10 genome. Sub libraries were combined. To visualize all nuclei simultaneously, scanpy (v1.10.4) was used to construct a KNN graph using the first 30 principal components. To cluster the cells, the Leiden algorithm was applied at a resolution of 1. Then, the harmony package (v0.0.10) was deployed to remove batch effects. Nuclei from the same organ (but different mice/sub libraries) were aggregated using Seurat (v5.1.0). Each organ was then clustered separately using scanpy and cell types and repair outcomes were assigned.

To assign cell types, each nuclei was scored for every possible cell type based on positive marker gene enrichment and negative marker absence in the CellMarker 2.0 database^69^. Cell type scores were then summed up across all cells in a cluster, and the top 3 cell types were evaluated manually before assigning a cell type to the cluster (**Table S4-9**).

### Assessing chromosome arm deletions and duplications using inferCNV

The inferCNV package (version 1.12.0) of the Trinity CTAT Project (https://github.com/broadinstitute/inferCNV) was used to infer copy number variations. In brief, gene expression measurements of unedited cells were subtracted from those of edited cells using base R and dplyr (version 1.1.4). The edited cells were then clustered based on their residual gene expression levels up to the target site (chr12:6,538,000 for *GAPDH* 3’ UTR targeting in human cells, chr2:122,153,000 for *B2m* 3’ UTR targeting in mouse cells). Subsequently, these data were summarized in a heatmap focusing on chromosome 12 using the packages ggplot2 (version 3.5.1) and ComplexHeatmap (version 2.20.0). Additionally, the mean residual gene expression of genes spanning from the start of chromosome 12 to the target site was calculated and compared to the mean residual gene expression of the following 7 Mb (chr12:6,538,000–13,638,000) as well as to the residual gene expression in unedited cells. This analysis was performed for each cell type individually to avoid cell-type-specific gene expression patterns.

### AAV cloning and production

AAV genome-containing plasmids (pAAV-scOUT-Son, pAAV-scOUT-B2m, pAAV-scOUT-NTC) were cloned by Gibson cloning starting from the AAV-Perturb seq backbone (Addgene #211502)^70^. These consisted of a HDR template (5’ homology arm, HDR barcode, 3’ homology arm), an sgRNA expression cassette and a Cre expression cassette with a T2A-dTomato or T2A-EGFP protein fused to a KASH domain to tag Cre expressing nuclei. AAVs harboring a non-targeting sgRNA were used as a control vector.

For murine LSK editing, AAV-6 serotype was produced targeting the *B2m* 3’ UTR, as described by the Platt lab^70^. AAVs were produced in HEK293T cells (Sigma-Aldrich, regularly tested for mycoplasma) and purified by iodixanol gradient centrifugation. In brief, HEK293T cells were expanded in DMEM (Merck) + 10% FBS (Merck) + 1% HEPES (Thermo Fisher Scientific). Then, 24 h before the beginning of AAV production, cells were seeded in 15 cm dishes (HuberLab) at a density of 0.6 M cells per ml and a total of 20 ml medium per dish. Cells were transiently transfected with 21 μg of an equal molar-ratio mix of the AAV genome, AAV6 plasmid (Addgene, #240485) and the adeno helper plasmid pAdDeltaF6 (Addgene, #112867) using polyethyleneimine max. At 48 h after transfection, the medium was replaced with fresh medium without FBS. The collected medium was mixed with 5× AAV precipitation buffer (400 g PEG 8000, 146.1 g NaCl in 1 l H_2_O) and kept at 4 °C. Then, 1 day later, the cells were mechanically dislodged and centrifuged at 800*g* for 15 min. The cell pellet was resuspended in 12 ml AAV lysis buffer (50 ml of 1 M TRIS-HCl (pH 8.5), 58.44 g NaCl, 5 ml of 2 M MgCl_2_ in 1 l) and flash-frozen in liquid nitrogen. The supernatant was mixed with 5× AAV precipitation buffer, added to the medium collected previously, incubated for 2 h at 4 °C and centrifuged at 3,000*g* for 1 h at 4 °C. The resulting pellet was resuspended in 3 ml AAV lysis buffer and added to the first cell pellet. The pellet was subjected to three freeze–thaw cycles and incubated with SAN (Merck) (50 U per 15 cm dish) for 1 h at 37 °C. After two centrifugation steps (saving the supernatant) at 3,000*g* for 15 min at 4 °C, 14.5 ml of the supernatant containing AAV particles was poured into an ultracentrifuge-ready tube (Beckman Coulter). Iodixanol gradients were prepared by sequential pipetting of the following iodixanol solutions: 9 ml (15%), 6 ml (25%), 5 ml (40%) and 5 ml (54%). Gradients were ultracentrifuged using the Beckman type 70 Ti rotor at 63,000 rpm for 2 h at 4 °C. To recover the AAV particles, the tubes were pierced at the bottom, 4 ml of gradient (mainly 54% phase) were allowed to pass through and discarded, and the next 3.5 ml (containing isolated AAV) were retained. To remove the iodixanol and concentrate the AAV, the solution was diluted with PBS + 10% glycerol and centrifuged through a 15 ml Amicon 100 kDa MWCO filter unit (Amicon) at 1,000*g* for 10 min. The dilution and centrifugation steps were repeated for three rounds. The resulting AAV solutions were aliquoted and flash-frozen in liquid nitrogen. The AAV particle concentration was determined by droplet digital PCR (ddPCR; BioRad). In brief, 5 μl of isolated AAVs were diluted 10× in water and treated with DNase I (NEB, M0303S) before preparing tenfold serial dilutions with ddPCR dilution buffer (Ultrapure Water with 2 ng μl^−1^ sheared salmon sperm DNA (Thermo Fisher Scientific, AM9680) and 0.05% Pluronic F-68 (Thermo Fisher Scientific, 24040032)). ddPCR reactions with primers targeting the KASH sequence (KASH_F, CTCAAGGGTGATCAGGGCAG; KASH_R, GCATGGGGTAGAAGGATCGG; KASH_probe, ACAGCTGCACCCAGGCCAAC) were performed with 5.5 μl of the diluted AAV template, 11 µl ddPCR supermix for probes (BioRad, 1863024), 0.9 µM of both primers and 0.25 µM probe in a total of 22 µl. Droplets were generated using the BioRad ddPCR apparatus according to the manufacturer’s indications. The amplification reaction was performed as follows: (1) 95 °C for 10 min; (2) 42 cycles of 95 °C for 30 s and 60 °C for 1 min (42 cycles); (3) 72 °C for 15 s; and finally (4) 98 °C for 10 min. Data were collected and analysed using the BioRad ddPCR apparatus to calculate the number of viral particles per µl.

For murine liver, skeletal muscle, lung and heart editing, AAV-8 and AAV-9 serotypes were produced by the viral vector facility of UZH targeting the *B2m* 3’ UTR.

For murine brain editing, AAV-PHP.eB serotype was produced by the viral vector facility of UZH targeting the *Son* 3’ UTR.

### *Ex vivo* editing of mouse HSPCs

AAV-6 targeting the B2m 3’ UTR was added to LSK cells at a titer of 50K viral genomes/cell. Three days later, GFP+ (=Cas9+) and dTomato+ (=infected) cells were sorted and single-cell gene expression as well as scOUT libraries were generated and analysed as described above.

### *In situ* editing of organs *in vivo*

For brain editing, all animal work was performed under the guidelines of the ETH Animal Welfare Office, the University Basel Veterinary Office and the Basel-Stadt Cantonal Veterinary Office under the license national numer 35693, BS3191. Mice were kept under specific-pathogen-free conditions on a standard light cycle. Female R26-LSL-Cas9-EGFP C57BL/6 mice aged 7-8 weeks were injected in the tail vein with 100 µl of AAV-PHP.eB at a titer of about 1 × 10^12^ viral genomes per mouse^71,72^.

For liver, skeletal muscle, heart and lung editing, animal experiments were performed according to protocols approved by the Kantonales Veterinäramt Zürich and in compliance with all relevant ethical regulations under the license national number 32794, ZH159/2020. For liver, skeletal muscle, heart and lung editing, female and male Cas9-EGFP C57BL/6 mice (Strain #:026175) aged between 8 and 51 weeks were injected via tail vein with 150 µl AAV-8 or AAV-9 at a titer of about 5 × 10^11^ viral genomes per mouse^71^. The mice were housed in a pathogen-free animal facility at the Institute of Pharmacology and Toxicology of the University of Zurich, and they were kept in a temperature- and humidity-controlled room on a 12-hour light-dark cycle. Mice were fed a standard laboratory chow (Kliba Nafag no. 3437 with 18.5% crude protein).

6 weeks after editing, tissues were extracted and flash frozen with liquid nitrogen. Nuclei were extracted from tissues using the Nuclei Extraction Buffer (Miltenyi Biotec, 130-128-024) according to the manufacturer’s instructions with small modifications (see modified version of nuclei extraction protocol). Extracted nuclei were then sorted using FACS for GFP^+^dTomato^+^ populations. From these populations, single-nucleus Evercode^TM^ whole transcriptome libraries as well as scOUT-parse libraries were generated.

### scOUT-parse library generation and analysis

The standard Evercode^TM^ whole transcriptome libraries (v2) were generated according to the manufacturer’s recommendations until recovery of first strand cDNA. During cDNA amplification, gene-specific primers for the B2m and Son loci were spiked into the cDNA amplification mix at 1% of the concentration of the cDNA amplification primers for the initial cDNA amplification. The targeted B2m or Son loci were amplified from 50 ng of cDNA using forward gene-specific primers in combination with a generic reverse sample index (SI)-PCR primer (20 cycles). The gene-specific primers contain a partial Illumina read 2 handle. The SI-PCR oligonucleotide anneals to the partial Illumina read 1 sequence at the 3′ end of the molecule which preserves the cell barcodes and UMI. After PCR1 unincorporated primers and large non-specific amplicons were removed using double-sided SPRI bead size selection (0.6X-0.8X). A subsequent second PCR was performed with a sample indexing primer (RPI-X) and a universal forward PCR primer to retain cell barcode and UMI information and index the samples (12 cycles). scOUT libraries were quantified using Quant-iT dsDAssay Kits (Thermo Fisher Scientific, Q33120) and their size distribution/molarity was assessed using D1000 ScreenTape on an Agilent TapeStation. scOUT libraries were sequenced on a NovaSeqX Plus and a NextSeq2000 Illumina system together with their parent Evercode^TM^ whole transcriptome libraries to ensure sufficient sequence diversity. The following cycle settings were used: 151 cycles for read 1, 151 cycles for read 2, and 8+8 cycles for the i7 and i5 sample index, respectively.

scOUT data was analyzed as above with taking into account the distinct library and barcoding structure. Barcode combinations and UMIs were extracted from the reads, and a whitelist is built out of these combinations. After salvaging barcodes, barcode IDs were mapped using the barcode manifest. The resulting scOUT data was then analyzed as described above using the CellMarker 2.0 database for cell type assignment^69^ (**Table S4-9**).

### Differential expression and gene-set enrichment analysis

To identify gene expression patterns correlated with repair outcome categories, cells were grouped according to their assigned repair outcomes. For each comparison of interest (e.g. HDR vs non-HDR, Insertion vs non-Insertion, NHEJ-Deletion vs non–NHEJ-Deletion) cells were divided into two groups based on the presence or absence of each given category in its alleles.

For instance, cells were labelled as “HDR” if they had at least one HDR allele, and as “non-HDR” otherwise. Differential expression analysis was performed between each pair of groups using the Wilcoxon rank-sum test implemented in Seurat (v5.1.0). Genes were considered differentially expressed if their FDR-adjusted p-value was below 0.1.

These lists of differentially expressed genes were then also used for Gene Set Enrichment Analysis (GSEA). GSEA was performed to assess enrichment of cell type-specific transcriptional signatures using the MSigDB C8 collection, biological process-specific transcriptional signatures using the MSigDB C5:GO:BP collection and KEGG pathway database, and molecular function-specific transcriptional signatures using the MSigDB C5:GO:MF collection. Genes were first ranked by their average log fold-change, and the ranked list was used as input for the GSEA function from the clusterProfiler package (v4.14.6). An FDR-adjusted p-value threshold of 0.05 was used to determine significance.

### Editing outcome enrichment

Unedited alleles were excluded from analysis. Editing outcomes were classified into mechanistically relevant categories (see editing outcome schematic). For each repair outcome class, we calculated its global (=bulk) average frequency across all cell types. We removed editing outcome categories that we could not clearly assign to a known repair pathway, and which were below a frequency threshold of 0.3% of all outcomes. We then quantified enrichment within each cell type by computing the fold change between cell-type-specific outcome frequency and the corresponding bulk average outcome frequency for every cell type-editing outcome pair. To answer the question whether editing-class usage is independent of cell type, or whether they are associated, we performed a G-test test for every editing class against the null hypothesis that the proportion of events in cell type X divided by the total events across all cell types is the same in every cell type. We used Benjamini-Hochberg FDR to correct for multiple testing.

### Co-occurrence bias

Unedited alleles were excluded from analyses. We also removed editing outcome categories that we could not clearly assign to a known repair pathway, and which were below a frequency threshold of 0.3% of all outcomes. Expected co-occurrence frequencies were calculated under the assumption of independent occurrence. The marginal frequency of each repair outcome, expressed as a percentage of total allele-level outcomes, was computed. Under a multinomial model analogous to Hardy–Weinberg genotype frequencies, the expected co-occurrence frequency of any non-homozygous outcome pair was calculated as twice the product of their marginal frequencies. Observed co-occurrence frequencies were counted symmetrically, meaning (A, B) and (B, A) were treated as equivalent. Frequencies were then normalized to sum to 100%. For each pair, the fold change of observed over expected frequency of co-occurrence was calculated and plotted in a correlation plot. To determine for each pair of outcomes whether it co-occurs more or less often than expected if the two events were independent, we calculated a χ² (chi-square) test statistic using observed and expected counts and performed the Benjamini-Hochberg false discovery rate (FDR) correction for multiple testing to arrive at an adjusted p-value for each pair.

### Indel-size differential

Unedited alleles, as well as any alleles that were not insertions or deletions (e.g. HDR, structural variants) were excluded from this analysis. The remaining editing alleles were binned by indel size. For each bin we calculated its overall frequency across all cells, as well as its frequency within every individual cell type. The indel-size differential for a given *cell type-indel size* pair was defined as the cell-type-specific frequency minus the corresponding bulk frequency, so positive values indicate enrichment and negative values indicate depletion. For each cell type, we first determined the total number of observed indel events. To control for variation in cell numbers, the bulk distribution was downsampled to the same total count using multinomial sampling proportional to its observed frequencies. This rarefaction step ensures that the comparison is not biased by unequal numbers of observed indels and reduces the risk of spurious zeros in sparsely sampled categories. We compared the rarefied cell type distribution against the downsampled bulk distribution the Kolmogorov–Smirnov statistic (maximal ECDF difference). To obtain empirical p-values, we used a permutation test under the null hypothesis that the cell type and bulk samples are drawn from the same underlying distribution. Specifically, the two rarefied samples were pooled, their labels were randomly permuted, and the test statistic was recalculated after splitting the shuffled sample into two groups of equal size. This procedure was repeated 2,000 times per comparison (unless otherwise specified). The permutation p-value was estimated as the fraction of permuted test statistics greater than or equal to the observed statistic. Because multiple cell types were tested against the same bulk distribution, resulting raw p-values were adjusted with Benjamini–Hochberg to control the false-discovery rate (FDR).

### Insertion sequence enrichment

Insertions were first stratified by the length of the inserted sequence. Within each length class we derived “bulk” nucleotide frequencies by tallying, at every insertion position, how often each base (A, C, G, T) appeared across the entire dataset. The same position-specific nucleotide frequencies were calculated independently for every cell type. Enrichment was then defined, for each base in each cell type, as the fold change between the cell-type–specific frequency and the corresponding global frequency for that insertion length and sequence position. To test for each cell type whether the distribution of insertion sequences is the same as the pooled (‘bulk’) distribution taken over all cell types we performed G-tests. The null distribution was obtained by Monte-Carlo permutation by drawing synthetic insertion vectors from a multinomial distribution with probabilities of occurrence in the bulk. Resulting raw p-values were adjusted with Benjamini–Hochberg to control the false-discovery rate (FDR).

### Data analysis

Schematics depicting experimental design were created with biorender.com.

GraphPad Prism 10 was used to generate several graphs.

Python (v3.11) was used for data analyses and statistical analyses and to generate several graphs.

R (v4.5) was used for data analyses and to generate several graphs.

## Supporting information

Key Resource Table

## Acknowledgements

We thank D.B. Kohn and Z.R. Garcia for their help with identifying suitable sentinel sgRNAs in human HSPCs. We also thank the ETH Genome Engineering and Measurement Laboratory, especially Z. Kontarakis and R. Ferreira Cássio, for providing Cas9 protein. We thank the ETHZ Flow Cytometry Core Facility for performing FACS on infected tissues and HSPCs. We acknowledge FGCZ for performing all next generation sequencing. We also acknowledge the Viral Vector Facility at UZH for AAV production. We thank K. Csorba and L. Kobel for assistance with supportive work and R. Viviani for administrative assistance and financial management. We thank the partial funding support provided by Cooperative Centers of Excellence in Hematology NIDDK Grant # DK106829 for allowing the purchase of HSPCs from the Fred Hutchinson Cancer Center at a greatly reduced fee. M.F.S. is supported by a Boehringer Ingelheim Fonds PhD fellowship (BIF 18919). J.E.C. is supported by the NOMIS Foundation, the Lotte and Adolf Hotz-Sprenger Stiftung, the Swiss National Science Foundation (project grants 310030_188858, 320030-227979, and 310030_201160), and the European Research Council (ERC) under the European Union’s Horizon 2020 research and innovation program (grant agreement No 855741, DDREAMM). This work was supported as a part of NCCR RNA & Disease, a National Centre of Competence (or Excellence) in Research, funded by the Swiss National Science Foundation (grant number 182880 and 205601). A.J.S. and R.J.P are supported by the Swiss National Science Foundation (51NF40-182895) and the Fickel Family Fund. Research in the Jackson laboratory is funded by Cancer Research UK (CRUK) Discovery Award DRCPBM\100005, CRUK core funding SEBINT-2024/100003 and European Research Council (ERC) Synergy Award Grant agreement no. 855741-DDREAMM-ERC-2019-SyG. S.L. and A.E. are supported by ERC Synergy Award 855741.

---

**Supplemental Figure 1.**
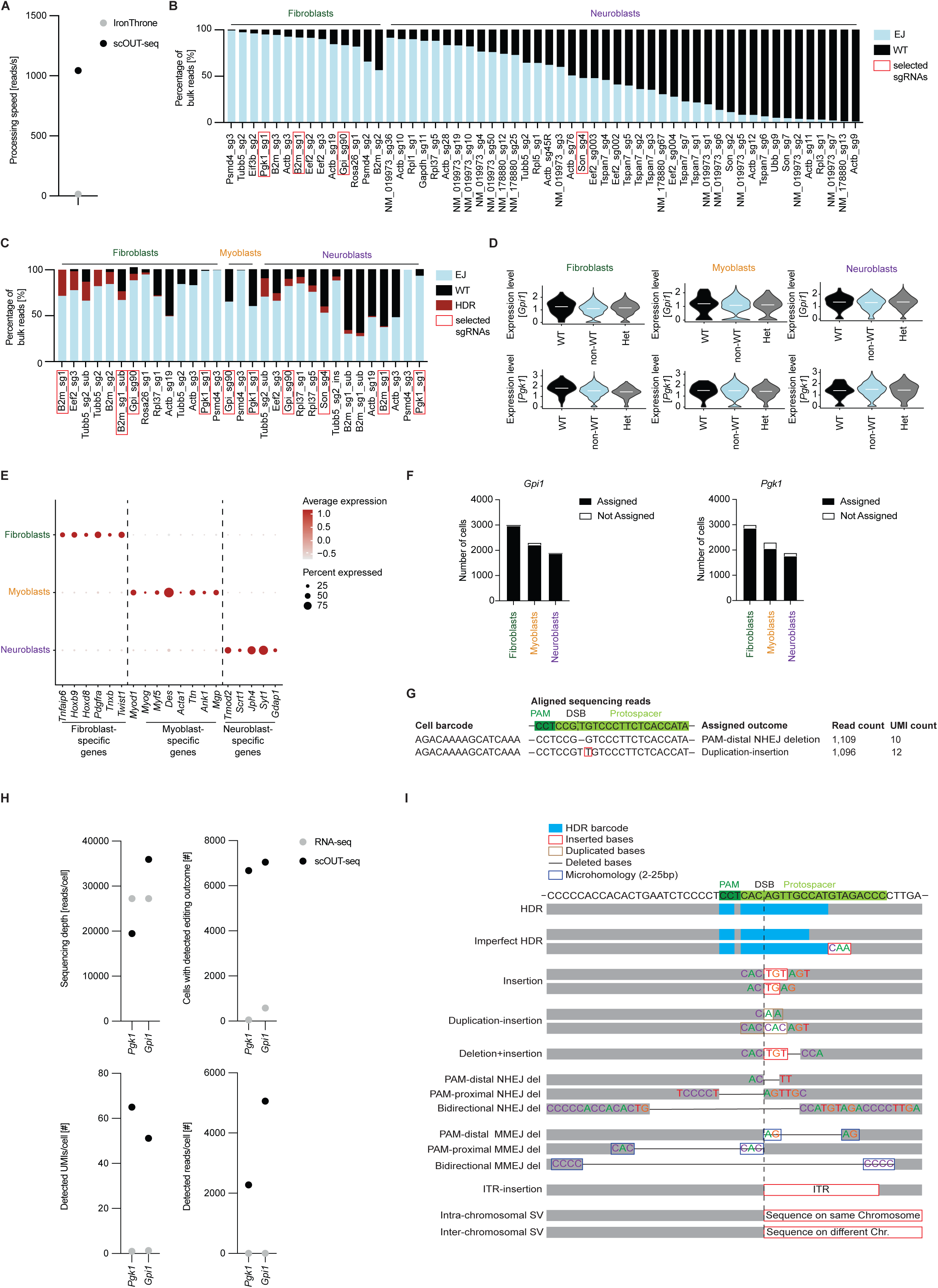
scOUT-seq accurately reflects bulk repair profiles, related to Figure 1. (A) Dot plot comparing processing speed between the IronThrone (GoT) and scOUT-MAP (scOUT-seq) pipelines. (B) Bar graph of editing efficiency without HDR templates of tested sentinel sgRNAs in mouse fibroblasts and neuroblasts. (C) Bar graph of editing efficiency with ssODN HDR templates of tested sentinel sgRNAs in mouse fibroblasts and neuroblasts. Insertion ssODN templates were used if not otherwise indicated by “sub” in the sgRNA name. Sub, substitution ssODN template. (D) *Gpi1* (*top*) and *Pgk1* (*bottom*) expression levels in cells with only unedited alleles (WT), only edited alleles (non-WT) and heterozygous cells (Het) in mouse fibroblast, myoblast and neuroblast cell lines. (E) Dot plot displaying expression of cell-line-specific genes across mouse fibroblast, myoblast and neuroblast cell lines. (F) Bar graphs showing the cell counts with or without assigned editing outcomes at *Gpi1* (*top*) and *Pgk1* (*bottom*) targets across indicated cell lines. (G) Example of scOUT-seq data showing biallelic editing outcomes in a single cell. PAM, protospacer-adjacent motif; DSB, double-strand break; UMI, unique molecular identifier. (H) Dot plots depicting sequencing depth (*top left*), number of cells with detected editing outcome (*top right*), detected UMIs per cell (*bottom left*) and detected reads per cell (*bottom right*) using scRNA-seq and scOUT-seq data. Data for the *Pgk1* and *Gpi1* loci are shown. (I) Schematic representation of editing outcome categories.

**Supplemental Figure 2.**
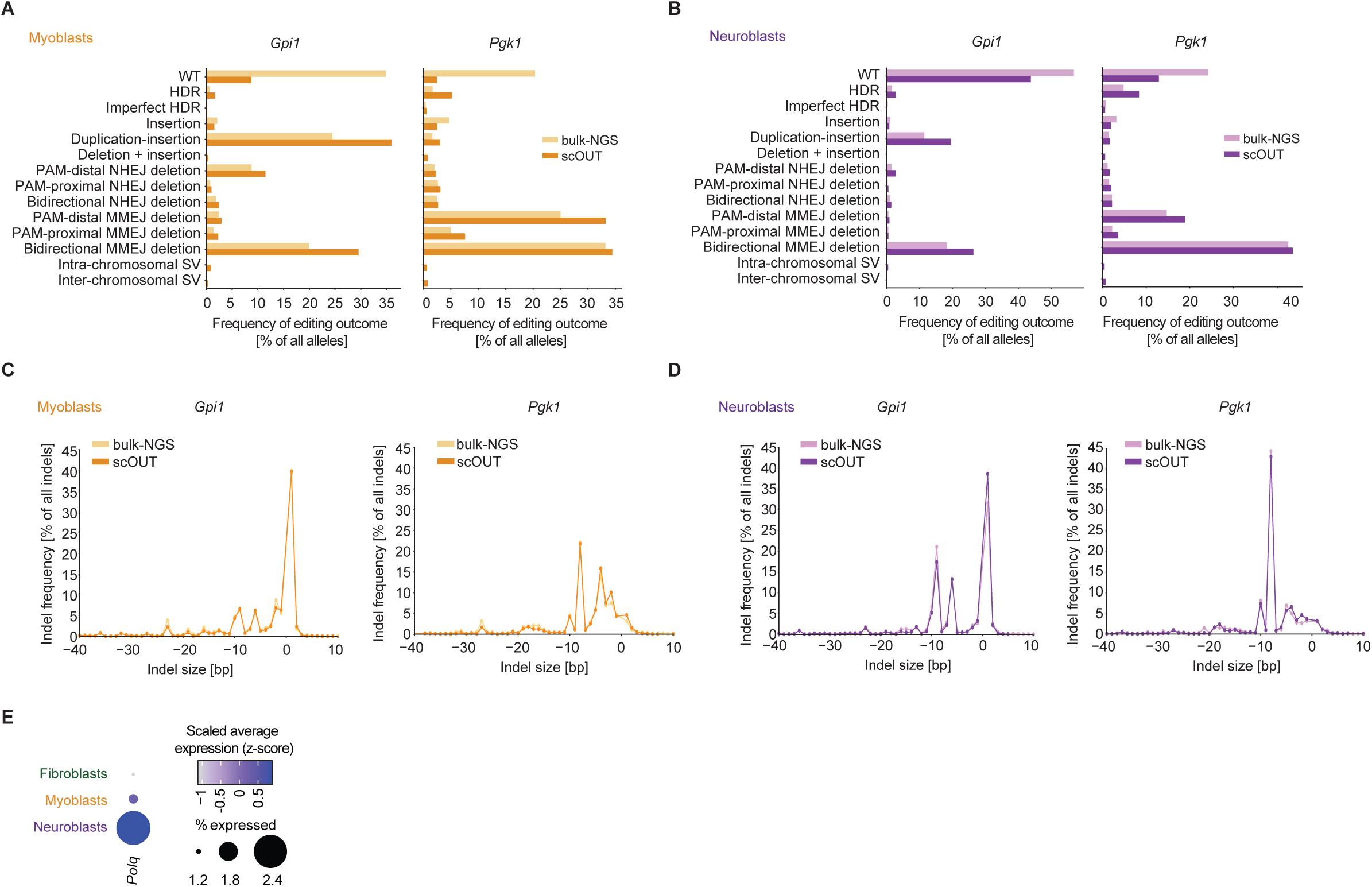
scOUT-seq accurately reflects bulk repair profiles, related to Figure 1. (A) Editing outcome frequencies at *Gpi1* (*left*) and *Pgk1* (*right*) loci in C2C12 mouse myoblast cells determined by bulk-NGS and scOUT-seq. (B) Editing outcome frequencies at *Gpi1* (*left*) and *Pgk1* (*right*) loci in Neuro-2a mouse neuroblast cells determined by bulk-NGS and scOUT-seq. (C) Indel-size frequencies at *Gpi1* (*left*) and *Pgk1* (*right*) loci in C2C12 mouse myoblast cells determined by bulk-NGS and scOUT-seq. (D) Indel-size frequencies at *Gpi1* (*left*) and *Pgk1* (*right*) loci in in Neuro-2a mouse neuroblast cells determined by bulk-NGS and scOUT-seq. (E) Dot plot displaying expression of *Polq* across mouse fibroblast, myoblast and neuroblast cell lines.

**Supplemental Figure 3.**
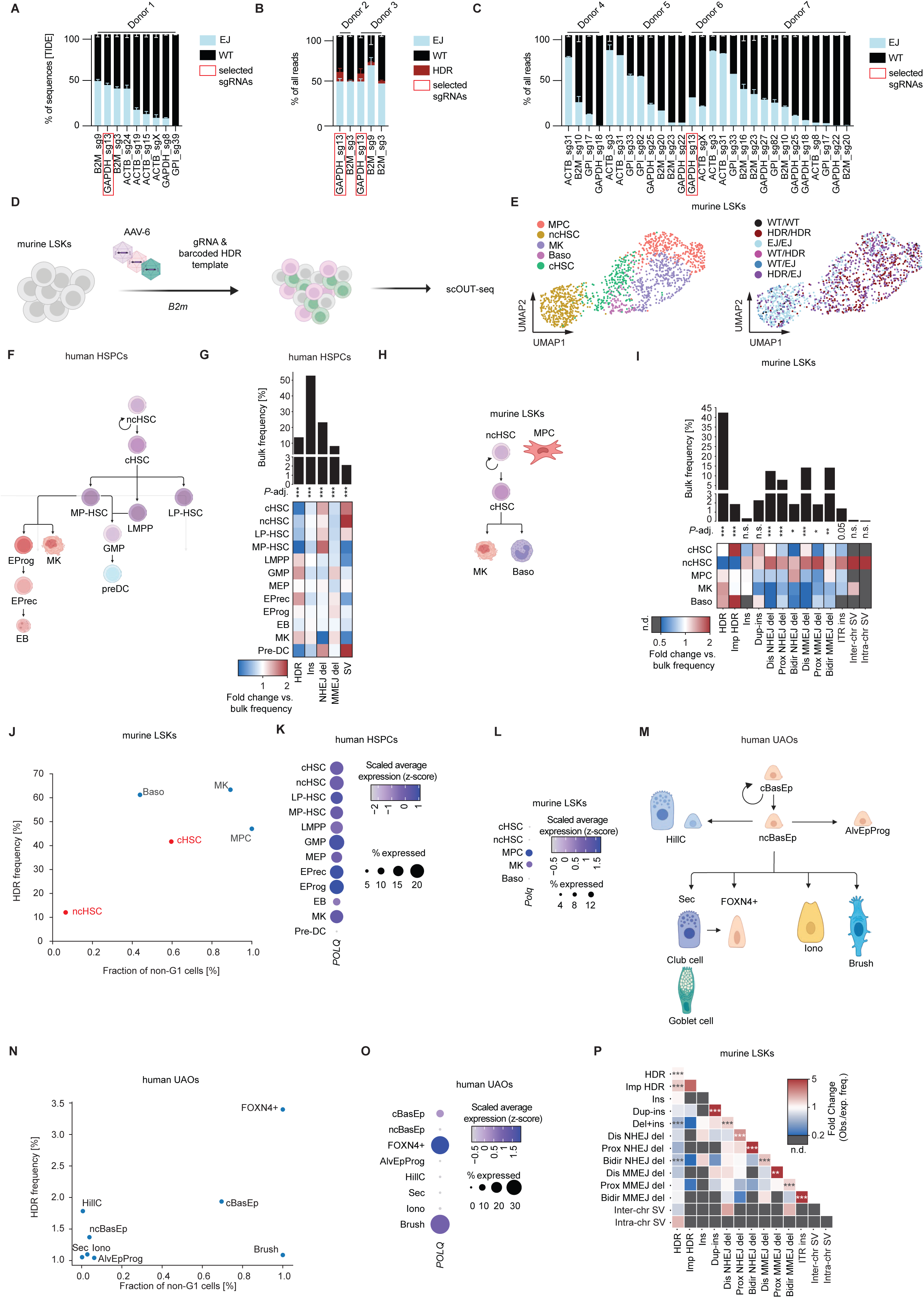
human CD34^+^ HSPCs and upper airway epithelial organoids, related to Figure 2. (A) Bar graph of editing efficiency without HDR templates of tested sentinel sgRNAs in human CD34^+^ HSPCs. (B) Bar graph of editing efficiency with ssODN HDR templates of tested sentinel sgRNAs in human CD34^+^ HSPCs. Substitution ssODN templates were used. (C) Bar graph of editing efficiency without HDR templates of additional tested sentinel sgRNAs in human CD34^+^ HSPCs. (D) Schematic of editing in murine Lin-Sca1+Kit- (LSK) cells using AAV-6. (E) UMAP plots displaying cell identities of 3,290 LSKs (*left*, color coded by cell line) and editing outcome per cell for 1,461 cells (*right*, color coded by editing outcome). MPC, mesenchymal progenitor cell; ncHSC, non-cycling hematopoietic stem cell; MK, megakaryocyte; Baso, basophil; cHSC, cycling HSC; WT, wild type; HDR, homology-directed repair; EJ, end-joining. (F) Lineage tree depicting the differentiation trajectories of human HSPC cell types. LMPP, lympho-myeloid primed progenitors; EProg, erythroid progenitors; LP-HSC, lymphoid-primed HSCs; EB, erythroblasts; GMP, granulocyte-monocyte progenitors; MEP, megakaryocyte-erythroid progenitors; cHSC, cycling HSCs; MK, megakaryocytes; ncHSC; non-cycling HSCs; EPrec, erythroid precursors; MP-HSC, myeloid-primed HSCs; Pre-DC, pre-dendritic cells (G) Bar graph displaying bulk frequencies of broad editing outcome categories in human CD34^+^ HSPCs (*top*). Heatmap of cell-type-specific fold-changes relative to bulk frequencies (*bottom*). FDR-adjusted p-values of G-tests determining difference of categories across cell lines are shown (***, *p*-adj. < 0.001). (H) Lineage tree depicting the differentiation trajectories of detected murine LSK cell types. MPC, mesenchymal progenitor cell; ncHSC, non-cycling hematopoietic stem cell; MK, megakaryocyte; Baso, basophil; cHSC, cycling HSC. (I) Bar graph of bulk frequencies of editing outcome categories in murine LSKs (*top*). Heatmap of cell-type-specific fold-changes relative to bulk frequencies. n.d., not detected. (*bottom*). FDR-adjusted p-values of G-tests determining difference of categories across cell types are shown (*, *p*-adj. < 0.05; **, *p*-adj. < 0.01; ***, *p*-adj. < 0.001; n.s., p-adj. > 0.05). (J) Scatter plot of fraction of non-G1 cells vs. HDR frequency per cell type. HSCs are colored in red. Cell types detected in murine LSKs are shown. (K) Dot plot of *POLQ* expression in cell types of human HSPCs. (L) Dot plot of *Polq* expression in cell types of murine LSKs. (M) Lineage tree depicting the differentiation trajectories of cell types detected in human UAOs. Sec, secretory cells; HillC, hillock cells; cBasEp, cycling basal epithelial cells; ncBasEp, non-cycling basal epithelial cells; Iono, ionocytes; Brush, brush cells, FOXN4^+^, FOXN4-positive cells; AlvEpProg, alveolar epithelial progenitors. (N) Scatter plot of fraction of non-G1 cells vs. HDR frequency per cell type. Cell types detected in human UAOs are shown. (O) Dot plot depicting *POLQ* expression in cell types of human UAOs. (P) Correlation plot of fold-changes between observed and expected frequency of co-occurrence (assuming independence of alleles) of editing outcome categories in murine LSK cells. n.d., not detected. FDR-adjusted p-values of a χ² (chi-square) test determining difference between observed and expected frequencies across cell types are shown (**, *p*-adj. < 0.01; ***, *p*-adj. < 0.001).

**Supplemental Figure 4.**
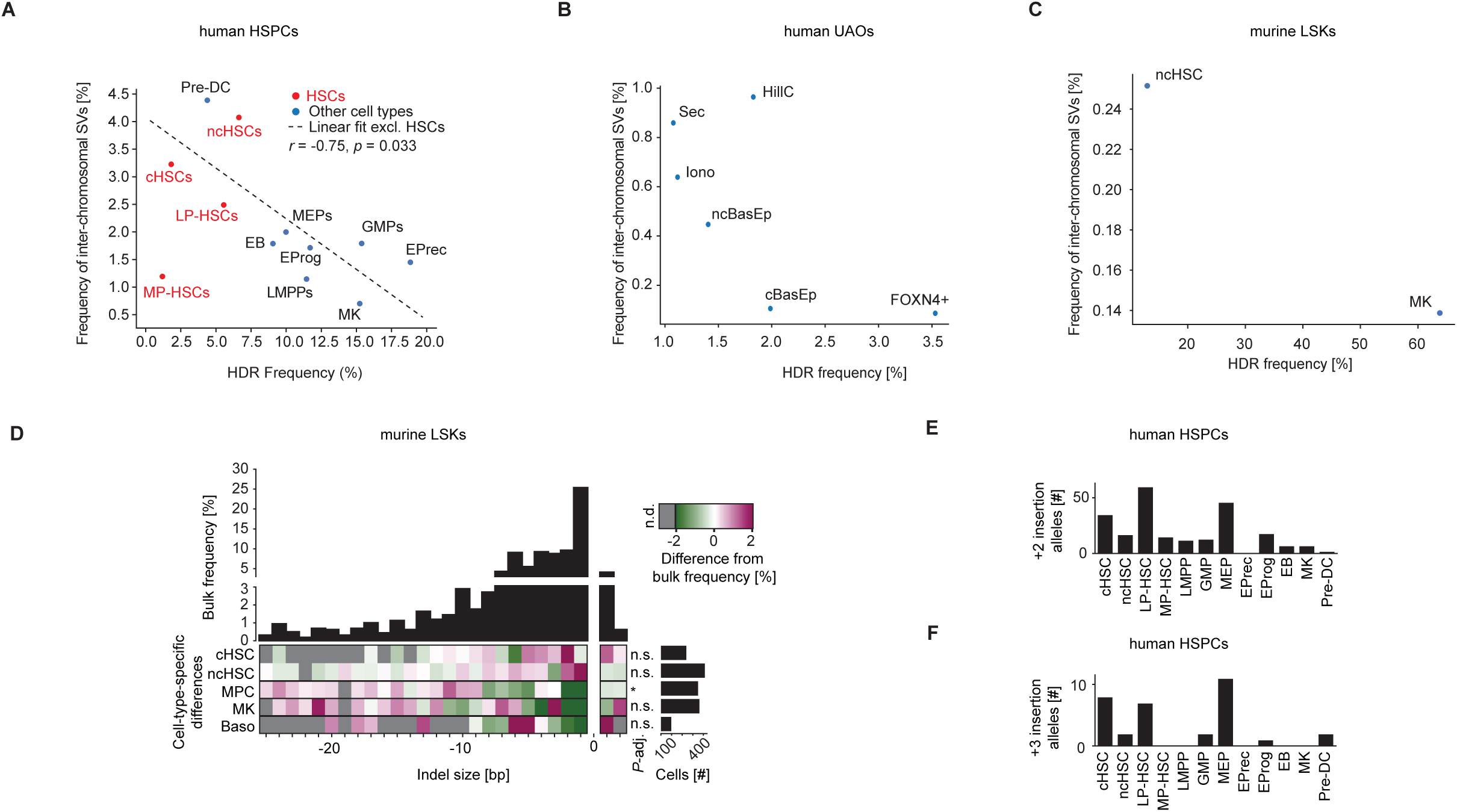
Editing outcomes in human CD34^+^ HSPCs and upper airway epithelial organoids, related to Figure 2 and 3. (A) Scatter plot of frequency of HDR vs. frequency of inter-chromosomal SVs per cell type. HSCs are colored in red. Cell types detected in human CD34^+^ HSPCs are shown. r, Pearson correlation coefficient; p, p-value. (B) Scatter plot of frequency of HDR vs. frequency of inter-chromosomal SVs per cell type. Cell types detected in human UAOs are shown. (C) Scatter plot of frequency of HDR vs. frequency of inter-chromosomal SVs per cell type. Cell types detected in murine LSKs are shown. (D) Histogram of bulk indel-size frequencies in murine LSK cells (*top*). Heatmap of percentage-point deviations in indel-size frequencies from the bulk across cell types in murine LSK cells with number of cells per cell type depicted in a bar graph. n.d., not detected. (*bottom*). FDR-adjusted p-values of G-tests determining difference of indel profile to bulk are shown (*, *p*-adj. < 0.05; n.s., p-adj. > 0.05). (E) Bar graph of number of +2 insertion alleles per cell type in human CD34^+^ HSPCs. (F) Bar graph of number of +3 insertion alleles per cell type in human CD34^+^ HSPCs.

**Supplemental Figure 5.**
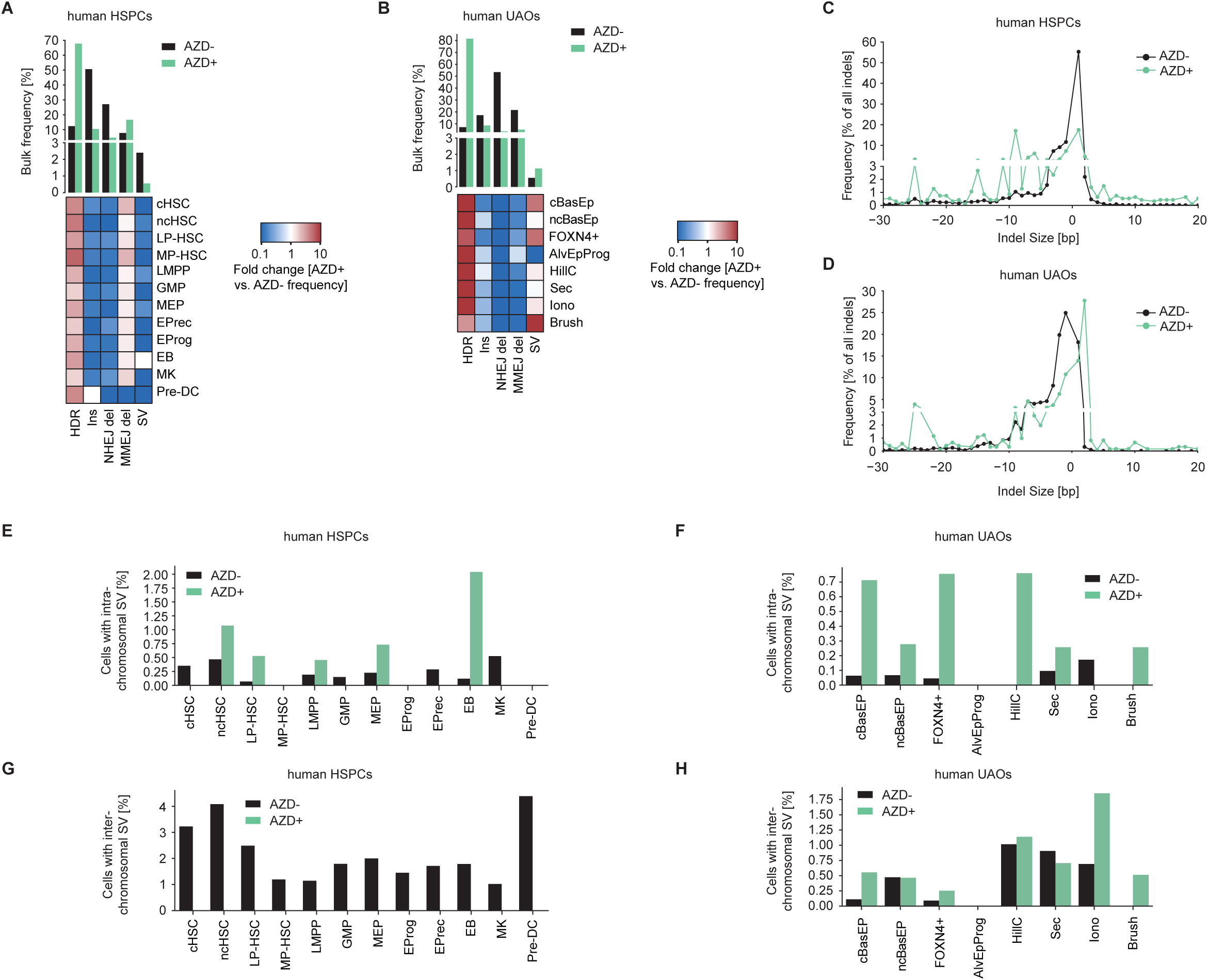
Cell-type-specific effects of DNA-PKcs inhibition during editing in human CD34^+^ HSPCs and upper airway epithelial organoids, related to Figure 4. (A) Bar graph showing bulk frequencies of broad editing outcome categories in AZD untreated (AZD-) and AZD treated (AZD+) human HSPCs (*top*). Heatmap of fold-changes in DNA-PKcs inhibited versus non-inhibited CD34*^+^* HSPCs across different cell types and editing outcome categories (*bottom*). (B) Bar graph showing bulk frequencies of broad editing outcome categories in AZD untreated (AZD-) and AZD treated (AZD+) human UAOs (*top*). Heatmap of fold-changes in DNA-PKcs inhibited versus non-inhibited human UAOs across different cell types and editing outcome categories (*bottom*). (C) Indel-size frequencies of DNA-PKcs inhibited versus non-inhibited CD34^+^ HSPCs. (D) Indel-size frequencies of DNA-PKcs inhibited versus non-inhibited human upper airway organoids (UAOs). (E) Bar graph depicting the fraction of cells within a cell type with intra-chromosomal SVs across different treatments in CD34^+^ HSPCs. NTC, non-targeting control; AZD+, edited with DNA-PKcs inhibition; AZD-, edited without DNA-PKcs inhibition. (F) Bar graph depicting the fraction of cells within a cell type with intra-chromosomal SVs across different treatments in human UAOs. NTC, non-targeting control; AZD+, edited with DNA-PKcs inhibition; AZD-, edited without DNA-PKcs inhibition. (G) Bar graph depicting the fraction of cells within a cell type with inter-chromosomal SVs across different treatments in CD34^+^ HSPCs. NTC, non-targeting control; AZD+, edited with DNA-PKcs inhibition; AZD-, edited without DNA-PKcs inhibition. (H) Bar graph depicting the fraction of cells within a cell type with inter-chromosomal SVs across different treatments in human UAOs. NTC, non-targeting control; AZD+, edited with DNA-PKcs inhibition; AZD-, edited without DNA-PKcs inhibition.

**Supplemental Figure 6.**
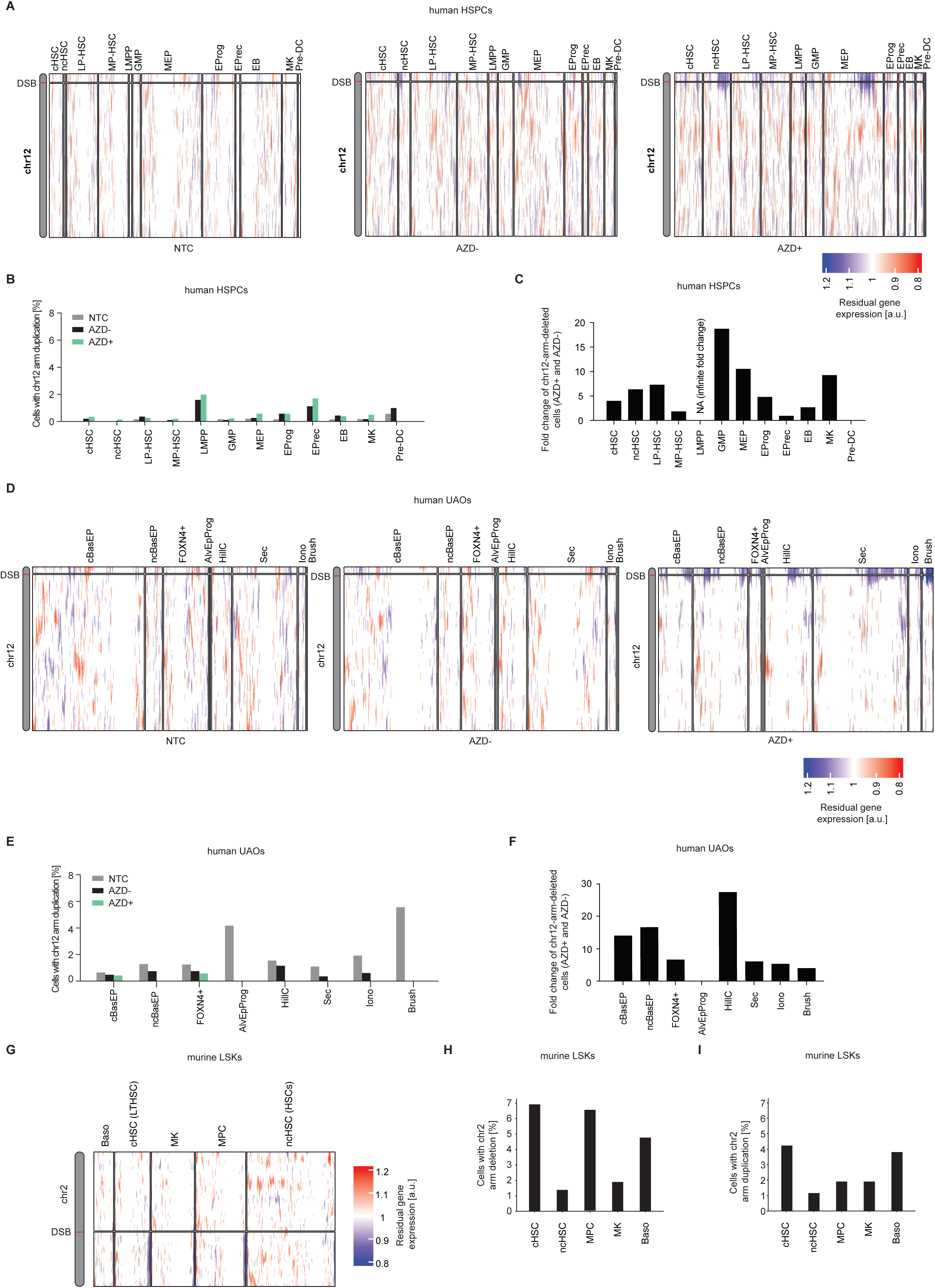
Cell-type-specific effects of DNA-PKcs inhibition during editing in human CD34^+^ HSPCs and upper airway epithelial organoids, related to Figure 4. (A) Heatmaps showing residual gene expression compared to unedited CD34^+^ HSPCs across chromosome 12 and grouped by cell types. The site of the DNA double strand break (DSB) induced by Cas9 is indicated. NTC, non-targeting control; AZD+, edited with DNA-PKcs inhibition, AZD-, edited without DNA-PKcs inhibition. (B) Bar graph depicting the fraction of cells within a cell type with chromosome 12 arm duplications across different treatments in CD34^+^ HSPCs. NTC, non-targeting control; AZD+, edited with DNA-PKcs inhibition; AZD-, edited without DNA-PKcs inhibition. (C) Bar graph depicting fold change of the number of cells with chromosome 12 arm deletions between AZD+ and AZD-human CD34^+^ HSPCs. AZD+, DNA-PKcs inhibition; AZD-, Cas9 edited. (D) Heatmaps showing residual gene expression compared to unedited human upper airway organoids across chromosome 12 and grouped by cell types. The site of DSB induced by Cas9 is indicated. NTC, non-targeting control; AZD+, edited with DNA-PKcs inhibition, AZD-, edited without DNA-PKcs inhibition. (E) Bar graph depicting the fraction of cells within indicated cell types with chromosome 12 arm duplications across different treatments in human UAOs. NTC, non-targeting control; AZD+, edited with DNA-PKcs inhibition; AZD^-^, edited without DNA-PKcs inhibition. (F) Bar graph depicting fold change of the number of cells with chromosome 12 arm deletions between AZD+ and AZD-human UAOs. AZD+, DNA-PKcs inhibition; AZD-, Cas9 edited. (G) Heatmap showing residual gene expression compared to unedited murine LSK cells across chromosome 2 and grouped by cell types. The site of the DSB induced by Cas9 is indicated. (H) Bar graph quantifying chromosome arm deletions across different cell types in murine LSK cells. (I) Bar graph quantifying chromosome arm duplications across different cell types in murine LSK cells.

**Supplemental Figure 7.**
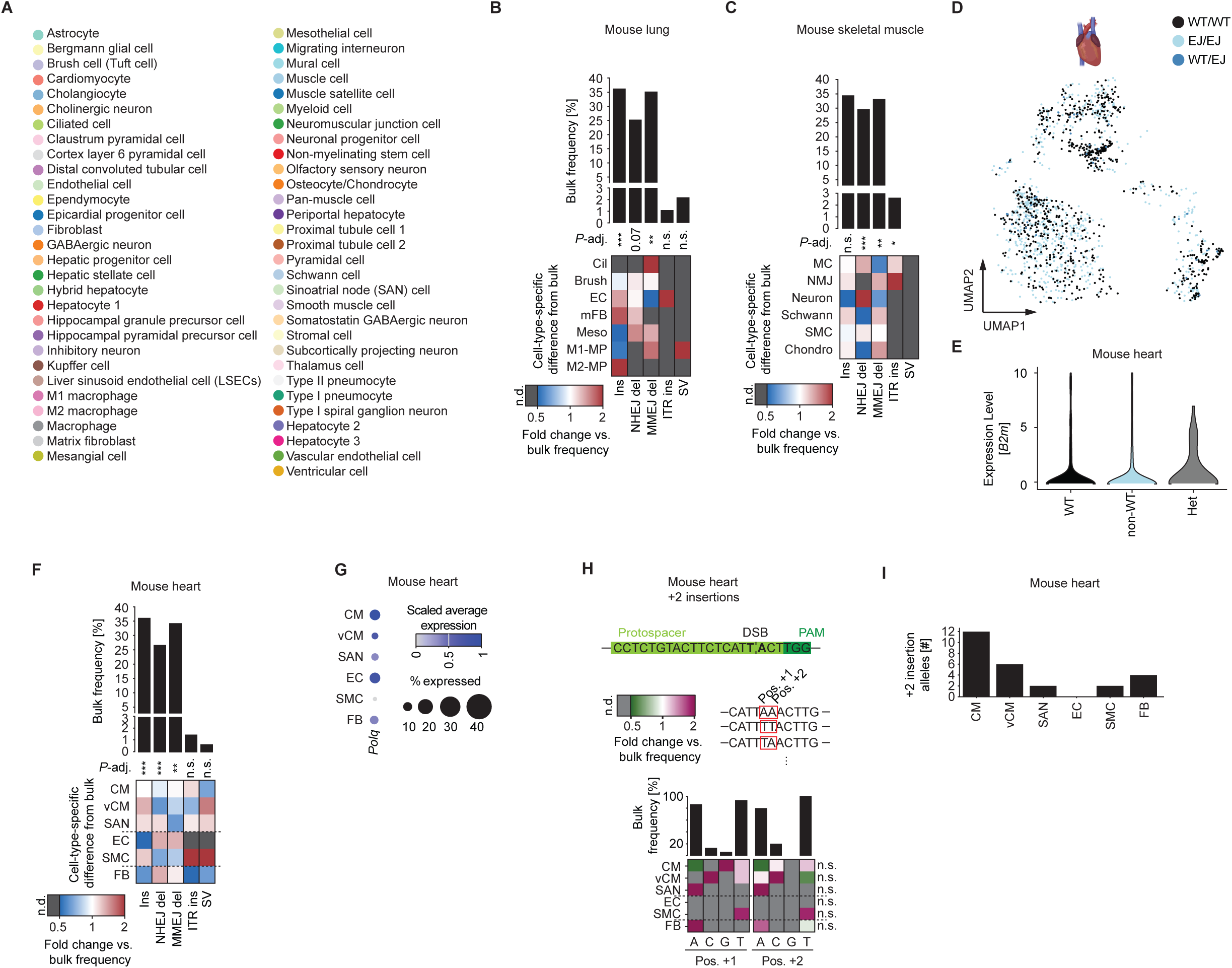
scOUT-seq across different tissues *in vivo*, related to Figure 5. (A) Legend for UMAP plot in **Figure 5C** indicating cell types. (B) Bar graph of bulk frequencies of broad editing outcome categories in mouse lung (*top*). Heatmap of cell-type-specific fold-changes relative to bulk frequencies. n.d., not detected. (*bottom*). FDR-adjusted p-values of G-tests determining difference of categories across cell lines are shown (**, *p*-adj. < 0.01; ***, *p*-adj. < 0.001; n.s., p-adj. > 0.05). (C) Bar graph of bulk frequencies of broad editing outcome categories in mouse skeletal muscle (*top*). Heatmap of cell-type-specific fold-changes relative to bulk frequencies. n.d., not detected. (*bottom*). FDR-adjusted p-values of G-tests determining difference of categories across cell lines are shown (*, *p*-adj. < 0.05; **, *p*-adj. < 0.01; ***, *p*-adj. < 0.001; n.s., p-adj. > 0.05). (D) UMAP plot of 1,582 mouse heart nuclei assigned with editing outcomes and colored by editing outcome category. (E) Normalized *B2m* expression levels in mouse heart nuclei with only unedited alleles (WT), only edited alleles (non-WT) and heterozygous alleles (Het) in mouse heart. (F) Bar graph of bulk frequencies of broad editing outcome categories in mouse heart (*top*). Heatmap of cell-type-specific fold-changes relative to bulk frequencies. n.d., not detected. (*bottom*). FDR-adjusted p-values of G-tests determining difference of categories across cell lines are shown (**, *p*-adj. < 0.01; ***, *p*-adj. < 0.001; n.s., p-adj. > 0.05). (G) Dot plot of *Polq* expression in cell types of mouse heart. (H) Insertion sequence composition of +2 insertions in mouse heart. Fold change compared to the bulk is plotted in a heatmap across cell types and insertion sequence positions. n.d., not detected. Bar graphs depict the bulk frequency of each base at each position. The sequence context of the *B2m* sgRNA, as well as examples of single base pair insertions, are depicted as a reference. FDR-adjusted p-values of G-tests determining difference of insertion sequence composition to bulk are shown (n.s., p-adj. > 0.05). CM, cardiomyocytes; vCM, ventral cardiomyocytes; SAN, sinoatrial node cells; EC, endothelial cells; SMC, smooth muscle cells; FB, fibroblasts. (I) Bar graph depicting frequency of +2 insertions in the different cell types detected in mouse heart.

**Supplemental Figure 8.**
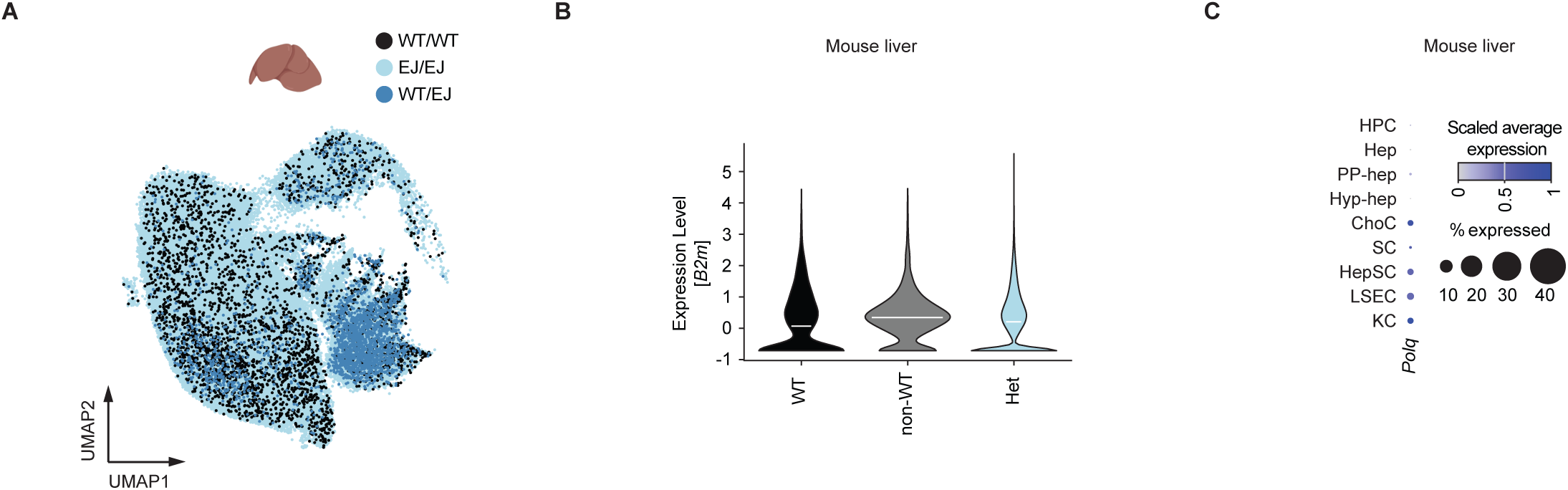
Mapping editing outcomes in mouse liver, related to Figure 6. (A) UMAP plot depicting 106,817 mouse liver nuclei assigned with editing outcomes and colored by editing outcome category. (B) Normalized *B2m* expression levels in nuclei with only unedited alleles (WT), only edited alleles (non-WT) and heterozygous alleles (Het) in mouse heart. (C) Dot plot depicting *Polq* expression in cell types of mouse liver. HPC, hepatocyte progenitor cells; Hep, hepatocytes; PP-hep, periportal hepatocytes; Hyb-hep, hybrid hepatocytes; ChoC, cholangiocytes; SC, stromal cells; HepSC, hepatic stellate cells; LSEC, liver sinusoidal endothelial cells; KC, Kupffer cells; ProxTC, proximal tubular cells.

**Supplemental Figure 9.**
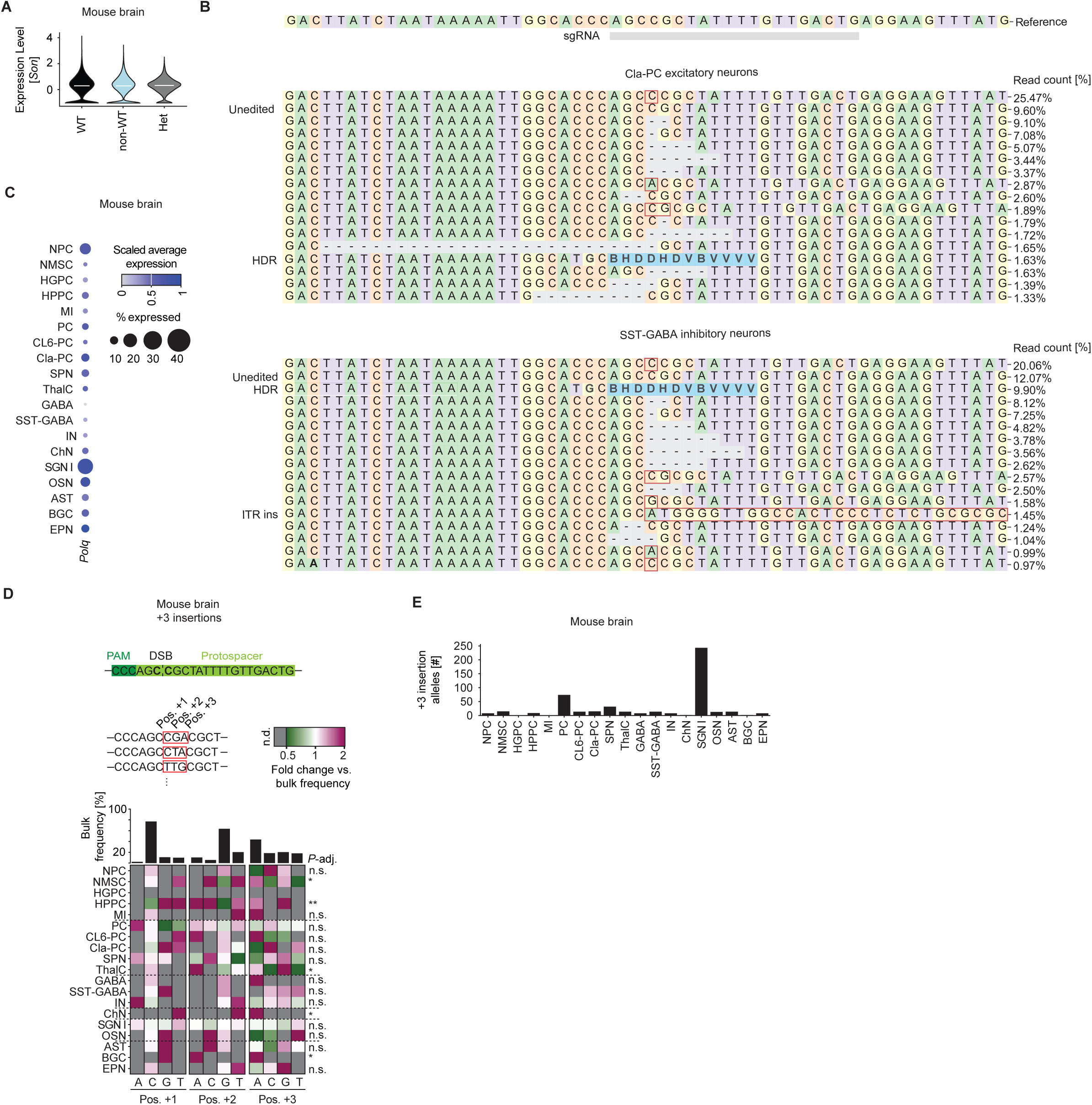
Mapping editing outcomes in mouse brain, related to Figure 7. (A) Normalized *Son* expression levels in cells with only unedited alleles (WT), only edited alleles (non-WT) and heterozygous nuclei (Het) in the mouse brain. (B) Allele plots of claustrum pyramidal cells (Cla-PC) and somatostatin-GABAergic neurons (SST-GABA). All HDR reads were combined to obtain their proportion of the total read count. (C) Dot plot depicting *Polq* expression in cell types of the mouse brain. (D) Insertion sequence composition of +3 insertions in the mouse brain. The fold change compared to the bulk is plotted across cell types and insertion sequence positions. n.d., not detected. Bar graphs depict the bulk frequency of each base at each position. FDR-adjusted p-values of G-tests determining difference of insertion sequence composition to bulk are shown (*, *p*-adj. < 0.05; **, *p*-adj. < 0.01; n.s., p-adj. > 0.05). (E) Bar graph depicting frequency of +3 insertions across cell types in the mouse brain.

